# Climate drives long-term landscape and rapid short-term promoter evolution in the Western Canaries lizard, *Gallotia galloti*

**DOI:** 10.1101/2024.12.18.628907

**Authors:** Edward Gilbert, Rodrigo Megía-Palma, Anamarija Žagar, Marta López-Darias, Miguel A. Carretero, Nina Serén, Graham S. Sellers, Pedro Beltran-Alvarez, Katharina C. Wollenberg Valero

## Abstract

Climate adaptation is caused by multiple mechanisms, including evolution of regulatory promoter sequences under broad environmental pressures, however, little is known about these processes in response to climate pressures in animals. Here, we examined spatial and temporal evolution of climate-related gene promoters in an insular species, the Western-Canaries lizard *Gallotia galloti*, across diverse environmental conditions over nine years. Outlier SNPs linked to elevation-sensitive environmental factors identified adaptively significant loci, including in aquaporin and Electron Transfer Flavoprotein Subunit Alpha (*ETFA*) gene promoters, demonstrating roles in local climate adaptation. Adaptive indexing mapped climate-driven genetic differentiation across the landscape, revealing local adaptation along an elevational gradient. Signatures of climate-related selection at the temporal scale showed specific promoters shaped by balancing selection supporting survival across environments, while others experienced positive selection, revealing a selective sweep of beneficial alleles. In addition, elevation-based ancestry groups of climate loci differed from the pattern observed in neutral loci, and temporal shifts in ancestry composition responded to preceding extreme weather events over the span of a single generation. Selection on specific promoter regions indicated adaptive evolution in transcription factor binding sites with the potential to enhance transcriptional flexibility, resulting in increased cellular stress tolerance. Our findings open new avenues into genomic mechanisms of resilience across spatially heterogeneous environments and how rapid evolution correlates with extreme weather events associated with climate change.

## Introduction

Anthropogenic climate change holds significant consequences for biomes and biodiversity (Román-Palacios & Wiens 2020; Foden et al. 2019). For example, it will push the thermal and hydric limits of animals (Rozen-Rechels et al. 2019; Greenberg & Palen 2021; Lertzman-Lepofsky et al. 2020; Buckley & Huey 2016), cause population-level disruption (Selwood et al. 2015; Kiritani 2013; Berggren et al. 2009), and be involved in future species extinctions (Urban 2015; Cahill et al. 2013; Sinervo et al. 2010). Simplistically, species facing climate change must adapt or risk extinction; a stark ultimatum shaping the future of life in the Anthropocene.

Climate adaptation is the process by which populations evolve to enhance their survival and reproduction in response to changing environmental conditions driven by climate (Frachon et al. 2018; Hoffmann & Sgrò 2011). In response to climate change, species can shift their distribution, elevation, and phenology (VanDerWal et al. 2013; Vitasse et al. 2021; Inouye 2022; Fuentes et al. 2024; Battisti et al. 2006; Sunday et al. 2012), change morphologically (Scriber 2020; Zeuss et al. 2014; Ficetola et al. 2016; Michel et al. 2017), demographically (Chevin et al. 2013), or alter their development and reproductive ecology (Jara et al. 2019; Perez et al. 2020; Servili et al. 2020; Llewelyn et al. 2018; Bodensteiner et al. 2023). Ectotherms rely on environmental heat sources to regulate their thermal physiology (Angilletta 2009) and may employ physiological and behavioural mechanisms to cope with shifting environments (Kearney et al. 2009; Bonebrake et al. 2014; Woods et al. 2015). Behavioural thermoregulation facilitates their optimal metabolism, reproduction, and overall life-history (Horváth et al. 2024; Beltrán et al. 2020; Stellatelli et al. 2024). However, in principle, if thermoregulation were completely efficient, physiological changes would be unnecessary (e.g. Bogert effect; Muñoz 2022). This suggests that a linear response to thermal heterogeneity might not be expected as would be observed in thermoconformers. Furthermore, while thermoregulation may buffer some impacts of environmental changes (at adult life stages), it may be insufficient to counteract the effects of a rapidly warming climate in the long term, or buffer vulnerable early life stages against extreme events (Sanger et al. 2018). Since physiological traits are ultimately encoded at the molecular level, and for populations to persist, change must occur as an evolutionary process (Cortés et al. 2020; Waldvogel, Feldmeyer, et al. 2020; Bodensteiner, Agudelo-Cantero, et al. 2021).

Climate adaptation at the molecular scale is often acknowledged, but rarely adequately addressed (Gienapp et al. 2008; Bodensteiner, Agudelo-Cantero, et al. 2021), although modern technologies have now begun to explore the role of epigenomics (McGuigan et al. 2021; McCaw et al. 2020), transcriptomics (Wang et al. 2021; Hamann et al. 2021; Rosso et al. 2024; Feugere et al. 2022), proteomics (Sun et al. 2022; Luo et al. 2024), and genomics (Saravanan et al. 2024; Wollenberg Valero et al. 2022; Turbek et al. 2023) in climate adaptation. The increasing availability of sequencing data has allowed scans for selection associated with traits or environmental parameters across genomes, a step change from studying a narrow range of predefined genes for adaptive evolution (Biswas & Akey 2006). This has uncovered an array of positively selected targets related to environmental adaptations (Rodríguez-Ramírez et al. 2023; Huerta-Sánchez et al. 2014; Stephan 2016), which contributes to understanding adaptive responses to environmental variation, potentially justifying a dedicated ontology of genes associated with environmental adaptation (Waldvogel, Schreiber, et al. 2020). After studies on the genomic basis of environmental adaptation and stress response accumulated, a pattern of extensive genomic reuse of specific genes with defined functions across diverse vertebrate clades and environmental gradients emerged (Rodríguez et al. 2017; Wollenberg Valero et al. 2014; Wollenberg Valero 2024; Wollenberg Valero et al. 2022). Deeper investigation of these genes, showing repeated use in climate adaptation, promises to further unravel the genomic mechanisms employed in adaptation to changing environmental parameters. Additionally, the roles of protein-coding genes and non-coding sequences, particularly *cis*-regulatory regions, in driving evolution and adaptation are key factors for understanding biological complexity (Taft et al. 2007; Planas & Serrat 2010; Castillo-Davis et al. 2004). The importance of non-coding sequences is underscored by the fact that many SNPs identified in genome-wide selection scans are located in non-coding regions (Uvarova et al. 2024; Zhang et al. 2021), such as promoters.

Promoter regions, which are DNA sequences upstream from the transcription start site of protein-coding genes, are critical for transcriptional control due to their role of providing binding sites for transcription factors and other regulatory elements (Andersson & Sandelin 2020).

Variation in promoter sequences can have significant downstream implications on gene expression and hence phenotypic differences (Hill et al. 2020; Ruan et al. 2020), therefore demonstrating an important source of selection driving evolution and adaptation (Young et al. 2015; Liang et al. 2008; Planas & Serrat 2010). As such, promoter sequences can drive adaptation to environmental stressors through both short-term transcriptional regulation, and long term environmental adaptation (López-Maury et al. 2008; Cases & Lorenzo 2005; Hoge et al. 2024). This mechanism has attracted little attention towards local adaptation in non-model vertebrates, but it can bridge the gap between genomics and transcriptomics, demonstrating evolutionary adaptation of expression changes, and not solely a snapshot of phenotypic plasticity.

Ectotherms are an ideal system for studying environmental adaptation in the context of climate change due to their reliance on external thermal and hydric gradients (Angilletta 2009; Siepielski et al. 2017; Baeckens et al. 2021; Cox & Cox 2015). Lizards, in particular, provide a useful model for testing climate related hypotheses, and have been the focus of several multi-taxa studies investigating their vulnerability, resilience, and relationship with climate (Rubalcaba et al. 2023; Jara et al. 2019; Lanna et al. 2022; Sinervo et al. 2010; Sinervo et al. 2024; Cosendey et al. 2022; Garcia-Porta et al. 2019).The Western-Canaries Lizard, *Gallotia galloti*, is a medium sized lacertid endemic to the islands of Tenerife and La Palma (Canary Islands), and the most abundant reptile inhabiting all environments of Tenerife. Tenerife’s diverse environments encompass an elevational gradient from sea level to the peak of El Teide stratovolcano over 3700 m, which is accompanied by a shift in vegetation structure and microclimate (Sperling et al. 2004). Humid coastal winds condensing on the high elevations, combined with elevation and orientation, support a generally more forested eco-zone in the North of the island, hosting laurel and pine forests (De Nascimento et al. 2009; Sperling et al. 2004). *G. galloti* occupies all bioclimatic zones on Tenerife, exploiting a broad niche across the island, with the subspecies *G. g. eisentrauti* present in the humid North and *G. g. galloti* in the South. *G. g. insulanagae* is only present in Roque de Fuera (offshore islets Roques de Anaga). Available phylogenetic analyses reveal two distinct mainland clades that do not align with subspecies distributions, suggesting that environmental factors may influence the species’ evolution (Richard & Thorpe 2001; Brown et al. 2016), making it an ideal system for studying climate adaptation.

This study assesses the spatial and temporal patterns of genetic variation and adaptation in *G. galloti* across an elevational gradient on Tenerife, examining how different environmental conditions between 2013 and 2022 have influenced the structure of 21 climate adaptation gene promoters. The coding sequences of these genes have previously been identified to harbour signatures of selection across 21 lacertid species including *G. galloti* (Garcia-Porta et al. 2019; Wollenberg Valero et al. 2022), shown transcriptomic response to heat in other vertebrates, or involved in water balance regulation. We hypothesised that genetic variation shows spatial and temporal structuring across topographical (elevation and orientation) induced environmental gradients and across time, with promoters exhibiting signatures of local adaptation at the extremes of an environmental gradient, and temporal changes aligning with extreme climate events.

## Materials & Methods

### Tenerife’s bioclimatic zones, sample collection and sequencing

Tenerife’s environment can be delineated into four main bioclimatic zones according to climate, altitude and orientation: hot, dry, low-lying coastal regions in the South (“Environment A”), intermediate temperate elevations (“Environment B”), cool high elevations (“Environment D”), and the humid North (“Environment C”) (Figure 1; Albaladejo et al. 2022; Algar & López-Darias 2016; De Nascimento et al. 2009). Sampling of *G. galloti* in Tenerife took place from 2013 to 2022, covering all Environments A–D, elevations, and all subspecies (Figure 1). DNA was extracted from 236 tissue samples using a standard salt-extraction method (Bruford et al. 1992) (sample details in Electronic Supplementary Materials). DNA quality and quantity were assessed by gel electrophoresis and quantified with a Qubit Fluorometer High Sensitivity (HS) assay.

**Figure 1.**
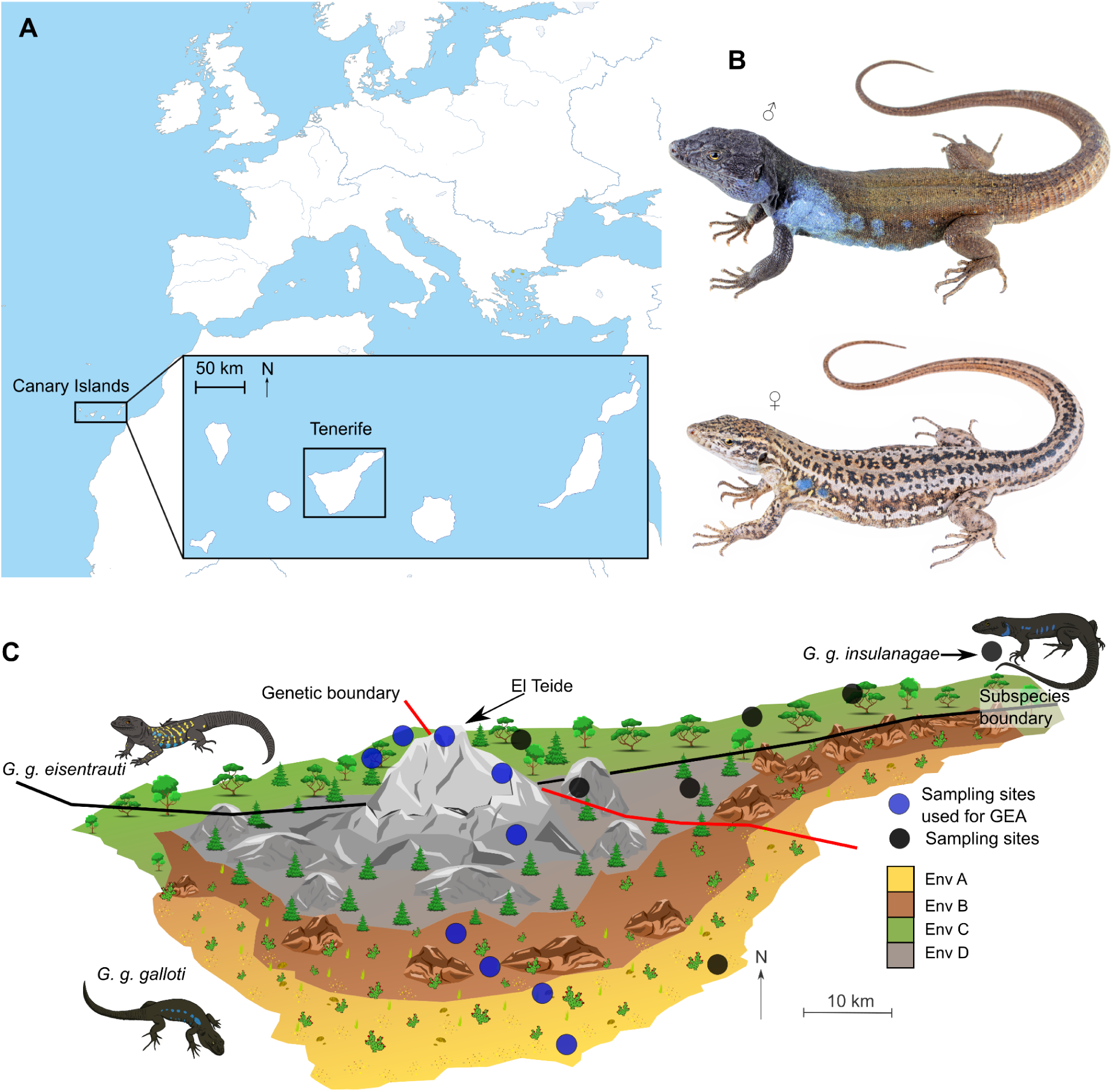
Sampling distribution of *Gallotia galloti* on Tenerife, Canary Islands. A) Canary Islands location map; off the coast of West Africa and position of Tenerife within the archipelago. B) male and female *G. g. galloti* from the South of Tenerife, photos reproduced with permission from Max Benito Smeele - ExSitu Project. C) schematic representation of Tenerife detailing the major environment types on the island (Env A - Env D) and approximate three-dimensional representation of the elevation, with the peak of El Teide stratovolcano labelled. The approximate distribution of *G. g. eisentrauti* and *G. g. galloti* are separated with a black line and phylogenetic clades with a red line (Thorpe et al. 1993; Richard & Thorpe 2001; Brown et al. 2006). *G. g. Insulanagae* only occurs on the offshore islet Roque de Fuera de Anaga. Sampling localities are shown with black circles, and North-South sampling localities used for the genotype-environment associations (GEA) are coloured in blue.

Fifteen genes relevant to climate and environmental adaptation across lacertid lizards were selected from Wollenberg Valero et al. (2022) and Garcia-Porta et al. (2019), subsetting to those with published links to heat stress response, and latitudinal adaptation in lacertids (details in Electronic Supplementary Material). Six additional genes from the Aquaporin (AQP) family relating to water balance were added from Zang et al. (2019). All these genes are referred to as Climate loci (CL). Mitochondrial Cytochrome B (*Cytb*), the mitochondrial control region (CR), and nuclear *RAG1* were included as putatively neutrally evolving genes (McClellan & McCracken 2001; Morales et al. 2015; Dias & Nery 2020). Bespoke primers were designed for each target region to maximise the fragment size, including an upstream promoter, the 5’ untranslated region (5’UTR) and the first exon (see primer design in Supplementary Methods). Each PCR amplicon was separately amplified for each sample (236 samples * 26 amplicons = 6,136 reactions), using LongAmp Taq Master Mix (New England Biolabs, M0287) according to the manufacturer’s guidelines (Supplementary Tables 1, 2). Amplicons were pooled per individual, balancing concentrations to ensure even representation (using the fluorescent band intensity for each amplicon as a proxy for concentration, quantified using gel electrophoresis images; details in Supplementary Methods). Pooled amplicons were cleaned using magnetic bead purification, quantified by Qubit, and prepared for sequencing using the Native Barcoding Kit 96 V14 and three MinION R10.4.1 flow cells. Sequencing ran on a MinION Mk1B for 72 hours per library (see details in Supplementary Methods).

Post sequencing, *Dorado* 0.6.2 (Oxford Nanopore Technologies) was used to basecall and demultiplex the pod5 outputs using the “Super Accurate Caller” (SUP) algorithm. *Fastp* (Chen 2023) was used to remove low quality reads (Q < 15), and the reads were mapped with the *map-ont* function using *minimap2* (Li 2017) to a reference library consisting of the target regions from the draft *G. galloti* genome (GCA_042478105.1; Serén et al. 2024). Coverage was calculated for each barcode across all the target amplicons using *samtools* (Danecek et al. 2021). *Bcftools* (Danecek et al. 2021) was used to perform haplotype and indel aware variant calling and generate respective consensus sequences. The VCF (variant call format genotype file) was filtered to exclude genotype calls informed by less than 10 reads, a mean genotype depth of 10 across all samples, SNPs (single nucleotide polymorphisms) when not all samples have sequence data, a quality value of 30, and with a minor allele frequency less than 5%. The VCF was converted into a numeric genotype matrix in R using the *vcfR* package (Knaus & Grünwald 2017). The consensus sequences were deconvoluted into alignments of all the samples for each target region using a custom script (Electronic Supplementary Materials).

### Population genetics

The population ancestry of all 236 samples from 17 localities over 9 years (7 sampled years) was calculated for CL and separately for NL. The *K*-value (number of ancestral populations) was estimated using the *snmf* function in *LEA* (Frichot & François 2015), with 10 repetitions and α of 500. Additional estimations of number of ancestry groups were performed with *ADMIXTURE* (Alexander et al. 2009), *dapc* from *adegenet* R package (Jombart & Ahmed 2011), and a scree-plot, yielding values between 3 and 6 for the NL, and 3 and 9 for the CL. A *K*-value of 3 was determined for the NL and 5 for the CL as the most reproducible values across the four methods. This was used to inform the *K*-value determined from population structure (using the NL, *K* = 3) for the LFMM in *LEA*.

Pairwise Nei F_st_ values were calculated for each year and environment combination using the *hierfstat* package in R (Goudet & Jombart 2022). Two year-environment combinations with less than 3 samples were removed (2016, Env B, Env D) and negative values were replaced with zero. Negative values represent more variation within a population than between populations, therefore were interpreted as no population differentiation (0). Half-matrix heatmaps were calculated for the CL and the NL. For the CL PCA, eigenvalues and eigenvectors from the VCF were calculated using PLINK (Purcell et al. 2007) and plotted in R with a custom script (Electronic Supplementary Materials).

### Climate data

Historical climate data of Tenerife was downloaded from WorldClim version 2.1 (Fick & Hijmans 2017) for 1970-2000, with bioclimatic variables (bioclim1-19), solar radiation, and elevation layers downloaded at the highest resolution available (30 arc-seconds, ∼1 km^2^). Annual minimum, maximum, and mean values were calculated for each layer from the monthly data.

Layers were cropped to Tenerife, and then tested for collinearity with the R package *corrplot* (Wei & Simko 2024), with variables removed when collinearity with another variable was > 0.9 (Supplementary Figure 1). Final selected variables were annual mean temperature, maximum temperature of the warmest month, annual precipitation, precipitation seasonality, mean solar radiation, and elevation. Elevation and annual mean temperature showed high correlation, however they were both retained for downstream analyses due to their pertinence in the study aims.

### Spatial analyses

To identify genotype-environment associations (GEAs), a subset of sequencing data was used (n = 174, 5 years, 10 localities) focusing on localities sampled over multiple years along an elevational transect through Tenerife (Figure 1). This excluded specific localities, such as Roques de Anaga and samples from the Northeastern clade, to reduce the influence of distinct genotypes from isolated environments. To detect associations between environmental variables and genetic variation, accounting for population structure, a latent factor mixed model (LFMM) was used, incorporating latent factors to reduce false positives. The numeric genotype matrix was generated from the VCF, formatted specifically for the *LEA* package (Frichot & François 2015). The K-value estimated using the *snmf* function in *LEA* and validated with other methods (described previously) was determined to be 3, using the NL as a proxy of population structure. Non-variable SNPs were removed from the genotype matrix, and the environmental data was consolidated into a single principal component representing 85% of variation, centred and scaled, for analysis. The LFMM was then run for 10 repetitions, and visualised using a dot Manhattan plot with a 95% significance threshold. Redundancy analyses (RDA) were conducted using the *vegan* package in R (Oksanen et al. 2024), integrating the genotype matrix, environmental variables, geographic coordinates, year of sampling, and population structure as predictors. Population structure was inferred through principal component analysis (PCA) using the *rda* functions without any predictors on the NL, with the first three PCs explaining 28.5% of the variance. Temporal variation was addressed by incorporating the year of sampling, spatial patterns were captured by geographic coordinates, and environmental variation was accounted for from the WorldClim variables. Variance partitioning comparing the statistical contribution of each of these predictors was performed through a set of partial redundancy analyses (pRDA).

This included a ‘full’ model in addition to a climate only, population structure only, geography only, and year only model. In the climate model, a model selection was carried out using the *ordistep* function in the *vegan* package, retaining annual mean temperature, elevation, annual precipitation, and mean solar radiation, as pertinent environmental variables. Once the final model was calculated, RDA scores were extracted at the sample level and at the SNP level, and plotted using biplots. For the SNP level analysis and to identify outlier SNPs, a threshold of 99% was applied using Mahalanobis distances following previous methods (Capblancq et al. 2018; Capblancq & Forester 2021). Outliers common to both simple (unconstrained) and partial (constrained) RDAs were classified as adaptively enriched SNPs, with results visualised spatially using an adaptive index following Capblancq & Forester (2021) (see details in Supplementary Materials).

### Temporal analyses

Allelic richness and nucleotide diversity (π) were assessed across years and environmental categories using *hierfstat* and *ape R* packages (Goudet & Jombart 2022; Paradis & Schliep 2019). They were fitted with general linear models (GLMs) with year-environment interactions to model trends. Average Tajima’s D values were calculated using the *pegas* R package (Paradis 2010) over the consensus sequences, and K-means clustering (K = 3) was applied to categorise temporal SNP trends by each environmental type.

Weather station data was obtained from AEMET (Agencia Estatal de Meteorologia) for four representative locations in Tenerife (Izana 28.3089, -16.4994 alt 2369 m, Los Rodeos 28.4667, -16.3167 alt 632 m, Tenerife Sur 28.05, -16.5667 alt 64 m, Santa Cruz de Tenerife 28.4682, -16.2546 alt 36 m) between 2012 and 2023 to estimate both contemporary trends and extreme weather events. Extreme hot and dry periods were calculated as the top 10% of maximum and mean temperatures per month, and bottom 10% of total precipitation per month, respectively, for each weather station. Extreme hot and dry periods were generated as count data for each year, and used alongside admixture proportions for CL in each year. Cross-correlation coefficient analysis was used to determine the lag between the climate events and changes in admixture proportion using the *ccf* function in base R, with a maximum lag of 6. Age structure from samples collected in 2022 was estimated based on recorded SVL (mm), using the equation *age = 0.060 x SVL - 1.013*, based on size-age relationships determined in (Castanet & Báez 1991), with no age structuring previously being detected across altitudes (Serén et al. 2023).

Permutation testing was applied to compare observed values against those expected by chance, with 1000 permutations and a max lag of 6, for each population (admixture group).

### Functional analyses

A sliding window of Tajima’s D was calculated across each consensus sequence using the *ape* (Paradis & Schliep 2019) and *pegas* (Paradis 2010) packages in R, which were visualised using a kernel density plot and a heatmap. The NL were plotted on a separate kernel density plot, but included in the heatmap. Sequences with high Tajima’s D value between -1000 and -700 bases upstream of the transcription start site (∼380 bp) were extracted from the promoters with the overall highest and lowest values of Tajima’s D (strongest signatures of selection), and scanned for potential transcription factor binding sites (TFBS) and motifs of interest through the JASPAR database (Rauluseviciute et al. 2024) with a 95% relative profile score threshold. A heatmap was used to visualise the mean relative score across each gene locus and identified TFBS. A sequence motif logo was generated from the position frequency matrix of nucleotides from the predicted sequences returned from JASPAR, filtering for the TFBS with the highest relative score (Ahr::Arnt complex).

## Results

### Spatiotemporal population genetics of *Gallotia galloti*

Admixture analysis identified spatial-temporal population structure and gene flow, revealing three ancestry groups in NL, two shared in the southwest, and one restricted to the North (Figure 2A-C). In the CL, five ancestry groups emerged, with less clear spatial structuring but showing broad North-South and northeastern-southwestern patterns (Figure 2A-C). Samples from 2022 displayed a large proportion of shared ancestry, while those from 2016-2021 were spatially segregated into Southern, Central, and Northern ancestry groups in the CL. Pairwise F_ST_ across years and environments further detailed genetic differentiation. Roque de Fuera de Anaga (RA) from 2014 showed marked differentiation in both climate and neutral loci (Figure 2B). Samples from 2022 were distinct from earlier samples, showing a shift towards overrepresentation of ancestry group CL1. Pairwise F_ST_ also indicated notable separation of environment C (distribution of *G. g. eisentrauti*) from other environments in the NL, particularly in 2021.

**Figure 2.**
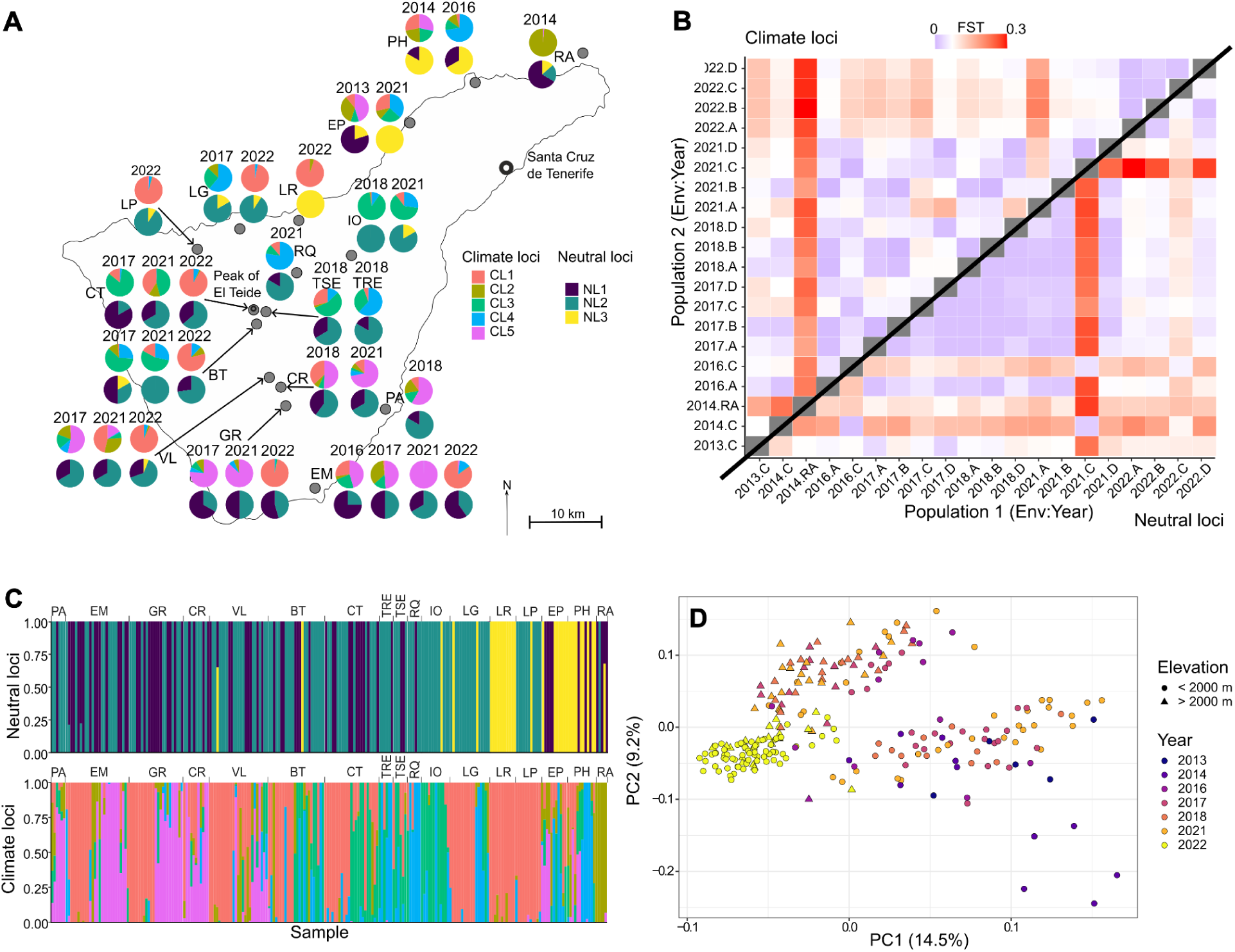
Spatio-temporal population statistics. A) Admixture proportions in climate loci (*K* = 5) and neutral loci (*K* = 3) from each sampling locality for each year in corresponding locations (see Electronic Supplementary Material for sampling locations). B) Pairwise F_ST_ calculations for each year-environment combination. The top left triangle shows the climate loci, and the bottom right triangle shows the neutral loci. C) Admixture proportions across climate loci and neutral loci for each sampling locality, corresponding to panel A. D) *PLINK*-Principal component analysis (PCA) from climate loci. Circles and triangles represent elevation less or greater than 2000 m, respectively. The year sampled is reflected as a colour gradient.

PCA of CL revealed genetic structuring across time and elevation (Figure 2D). No separation between *G. g. galloti* and *G. g. eisentrauti* appeared, however two clusters were identified separating high (2000 - 3700 m) and low (0 - 2000 m) elevation populations along the tree limit on the PCA2 axis (9.2%). A temporal gradient emerged along the PCA1 axis (14.5%), with 2022 clustering tightly in negative PCA space, converging from older samples (2013-2021) in positive values across the two elevational bands, suggesting more homogenous CL over time.

### Spatial changes in promoter sequences

The LFMM identified 23 SNPs potentially under selection for spatial environmental variables in PC1, explaining 85% of the variance (Figure 3A). A strong association is observed for *AQP11* (*AQP11*_1809), along with SNPs in *PHB1*, and several other AQP genes. Numerous SNPs associated with the *ETFA* gene were detected as outliers, corroborated by the RDA. These SNPs were located in areas of positive Tajima’s D, rather than the small section of negative Tajima’s D. The RDA showed the axes RDA1 and RDA2 explained a cumulative 3.34% of variation. Samples from the same environment clustered together, with Environments A and B forming one group and aligning with annual mean temperature, while high elevation factors associated with Environment D. Environment B and C separated along RDA2, not aligned with any environmental variables, suggesting additional unmeasured factors. Only one SNP (*UGP2*_580) was exclusive to the LFMM, indicating that outlier loci are robust across different statistical approaches. Both LFMM and RDA analyses emphasise the *ETFA* and AQP gene regions, although the specific SNPs varied slightly. For instance, 6 out of 7 AQP SNPs from the adaptively enriched pRDA were also detected in the LFMM, with the other identified in the unconstrained RDA (*AQP4*, 777). Outlier SNPs visualised in pRDA showed 97 statistically significant outliers based on Mahalanobis distances, including 40 associated with *ETFA* (Figure 3D). Nine SNPs were unique to southern localities (Environments A, B; labelled in brown text on Figure 3D). In the correlation-based scaling, *ETFA* outliers showed high positive correlation with solar radiation, elevation, and precipitation, while *AQP11*_1809 correlated strongly with precipitation. These outliers were inversely correlated with annual mean temperature. Outliers specific to southern localities showed no strong correlation with measured environmental variables, and “*HSPA5*_1984,” “*SLC25A33*_995,” and “*ODC1*_2653” showed weak association with mean annual temperature. Gene ontology (GO) analysis revealed that outlier SNPs are enriched in terms linked to mitochondrial energy production, enzymatic processes, and ROS production, with Aquaporin genes involved in membrane transport and immune response (Figure 3B). Some genes were also associated with cellular stress and developmental processes.

**Figure 3.**
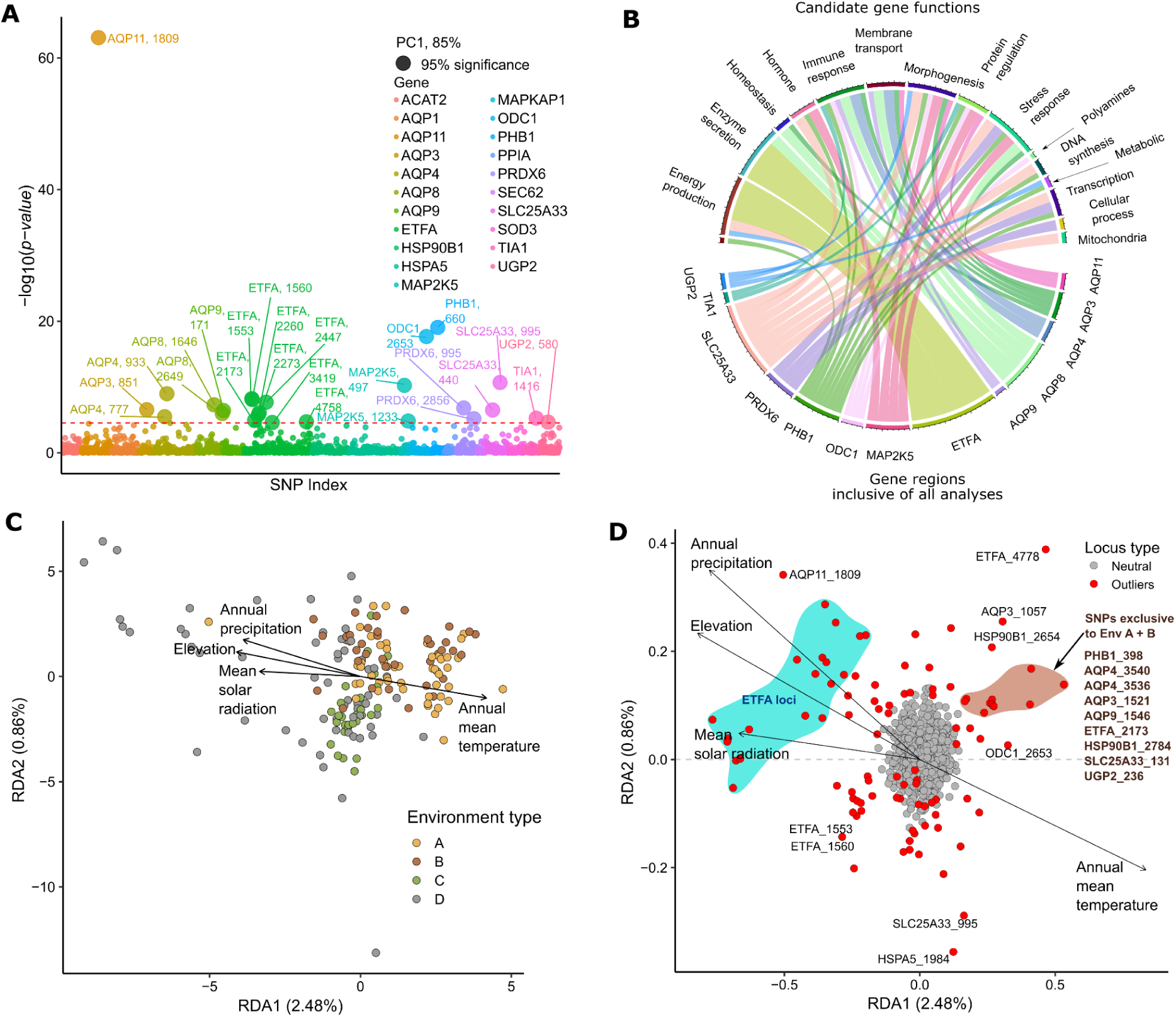
Genotype-Environment Associations (GEA). A) Latent Factor Mixed Model (LFMM) results derived from spatial-temporal genotype matrix and environmental variables consolidated into a single principal component (85%), with *p*-values plotted on a logarithmic axis (Log_10_). Coloured points represent SNPs associated with different gene regions. SNP outliers passing the 95% significance threshold are increased in size and labelled with the associated gene, and the SNP position. B) Candidate functions of each gene that occurred as an outlier in all GEA’s. The size of the chord linking the genes to the function is representative of the number of outlier SNPs detected in that promoter. Candidate gene functions are summaries of Gene Ontology (GO) terms to improve readability (see Electronic Supplementary Materials). C) Sample level partial redundancy analysis (pRDA). Samples are projected with the environmental variables along the first two RDA axes. Different colours represent samples from different environment types. D) SNP-level pRDA. The SNPs are projected with the environmental variables along the first two RDA axes. Red points indicate statistically significant outliers, with those furthest from the centroid labelled. Non-statistically significant SNPs are coloured grey. The blue cloud represents multiple statistically significant SNPs associated with the *ETFA* promoter. The brown cloud represents multiple statistically significant SNPs that are exclusive to the Southern part of the island (Environments A and B). These SNPs are labelled in brown font.

Further filtering based on inclusion in constrained and unconstrained RDA models isolated 83 SNPs (66%) for an “adaptively enriched RDA”, which explained 7.36% of data variance.

Retained outliers included *ETFA* promoter sites and SNPs exclusive to southern localities (Figure 4A). The same relationship between SNPs and environmental variables was observed as in Figure 3D. RDA1 explained 5.29% of variance (Figure 4A), showing allele frequencies positively associated with elevation, precipitation, and solar radiation, and negatively associated with annual mean temperature. When mapped, the adaptive index from RDA1 revealed an elevation-aligned gradient distinguishing high elevations at the island centre from coastal regions (Figure 4B, left). The RDA2 axis, explaining only 2.07% of variance, presented a higher adaptive index for another set of SNPs specific to mostly coastal areas (Figure 4B, right).

**Figure 4.**
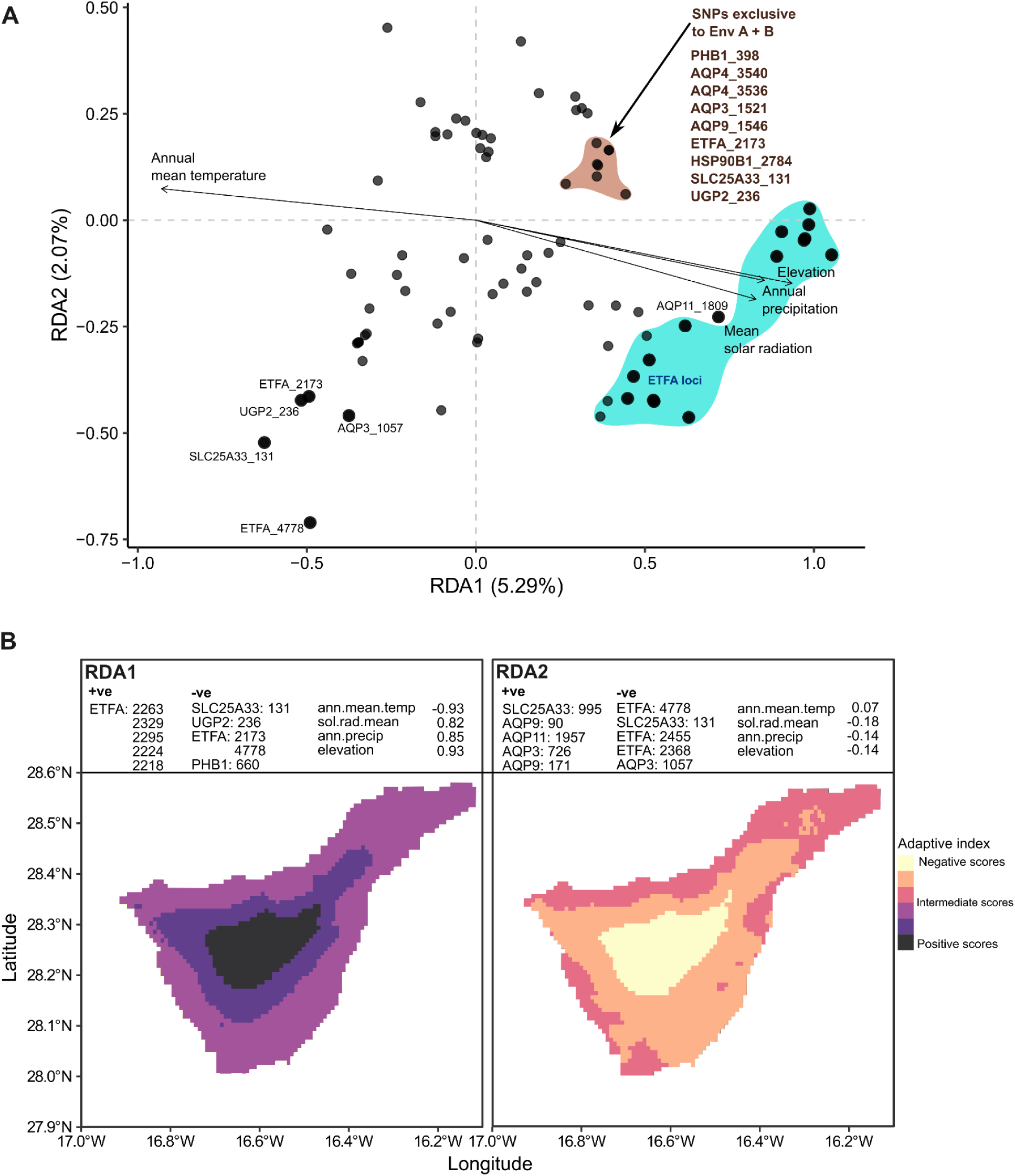
Adaptively enriched pRDA and adaptive index across landscape. A) SNP-level GEA using adaptively enriched pRDA. The SNPs are projected with the environmental variables along the first two RDA axes. All points are outliers determined from the previous pRDAs, with those furthest from the centroid labelled. The blue cloud represents multiple statistically significant SNPs associated with the *ETFA* loci. The brown cloud represents multiple statistically significant SNPs that are exclusive to the Southern part of the island (Environments A and B). B) Adaptive landscape of *Gallotia galloti* on Tenerife, using adaptively enriched genetic space to project adaptive indices across the species range. Prediction from RDA1 is on the left, and RDA2 is on the right. Values represent predicted adaptation based on environmental values in that pixel and SNP outlier frequency. Environmental variable loadings for each RDA are included, alongside the SNP associations with the most positive and negative associations in respective RDA axes.

### Temporal changes in promoter sequences

Temporal changes explained 9% of genetic variance (47% of explained variance; Supplementary Table 3). Nucleotide diversity remained constant across years and environments (Figure 5A, Supplementary Tables 6 and 7). ANOVA showed significant differences in allelic richness by environment types (*p* < 0.001), years (*p* < 0.001), and year-environment interactions (*p* = 0.044) (Supplementary Tables 4 and 5). Mean allelic richness was nearly equivalent in Environment A and C, while B and D had higher averages (Figure 5B). Environment B showed a significant interaction with years (*p* = 0.021) (Supplementary Table 4, and see the steeper slope in Figure 5B).

**Figure 5.**
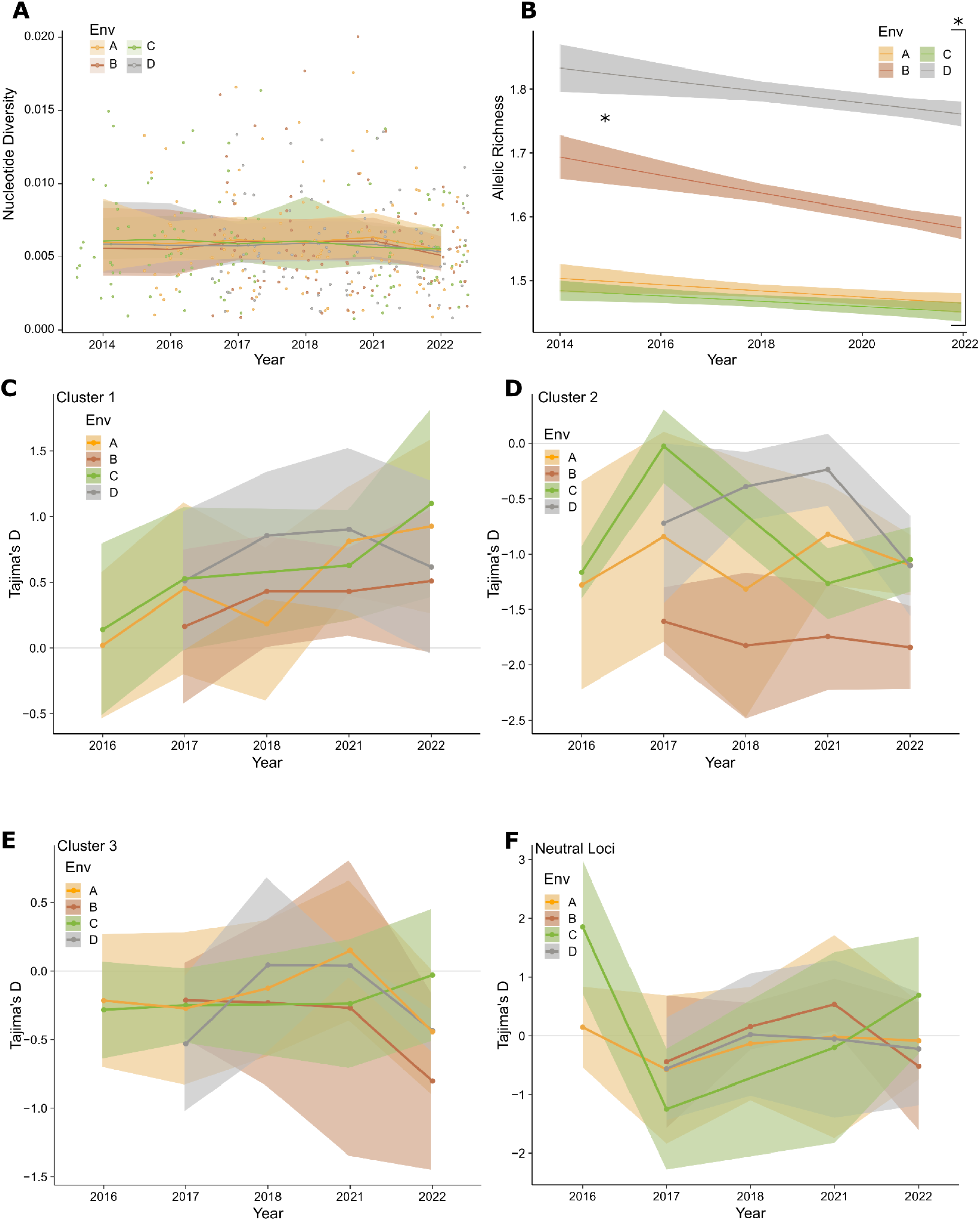
A) Mean nucleotide diversity for each major environment across different years. B) Mean allelic richness across all loci for each major environmental type, across different years. A) and B) are predictive GLM’s with standard error confidence intervals. C) - F) show K-means clusters of mean Tajima’s D values ± standard deviation (SD) from each year. Cluster 1 (C) shows genes with positive Tajima’s D values grouped by environment type, (D) cluster 2 shows genes with negative Tajima’s D values grouped by environment type, cluster 3 (E) shows genes with Tajima’s D values approximating zero grouped by environment type, and F) shows the neutral loci grouped by environment type. The grey horizontal line denotes D=0.

Clustering analysis of CL identified three distinct clusters based on Tajima’s D values and environmental patterns. Cluster 1, characterised by increasing Tajima’s D values over time, included genes such as Aquaporins (*AQP1*, *AQP11*), *ETFA*, *PRDX6*, and *ODC1* (Supplementary Figure 5A). This cluster shows evidence of increased balancing selection, particularly in environments A, B, and D, though AQP genes (*AQP8* and *AQP11*) experienced a sudden drop in Tajima’s D during 2021-2022, which is not consistent across all environments (Figure 5C). Cluster 2, marked by negative Tajima’s D values over time, contains genes including *HSP90B1* and *SLC25A33* (Supplementary Figure 5B), with a significant drop in Tajima’s D observed for *ACAT2*, suggesting these genes are under purifying selection, particularly in environments A and B (Figure 5D). Cluster 3 includes genes with neutral Tajima’s D values, such as *AQP4*, *HSPA5*, *MAP2K5*, *SOD3*, and *TIA1*, which are associated with stress responses (Supplementary Figure 5C). This cluster, as well as the NL remain relatively stable across years and environments, with no significant selection trends (Figures 5E, 5F).

Climate data from four weather stations identified 2017 and 2023 as particularly hot and dry years (the latter irrelevant to the genetics data as it post-dates the sampling period; Figures 6A, 6B). Changes in the ancestry group showed CL5 peaking in 2014, CL4 in 2016, CL2 dominating from 2018 to 2020, and a sharp rise in CL1 in 2022, indicating rapid haplotype turnover (Figure 6C). CL3 proportions remained consistent across years. Cross-correlation analysis linked CL1 positively with hot, dry periods at a 5-year lag (2017 weather with 2022 admixture), while CL2, CL3, and CL4 showed negative trends, strongest at a 5-year lag (Figure 6D). CL5’s correlation decreased with lag, reaching zero at 5 years. Permutation tests showed significance for CL4 and CL5 (p = 0.02 and p = 0.01, respectively; Supplementary Table 9).

**Figure 6.**
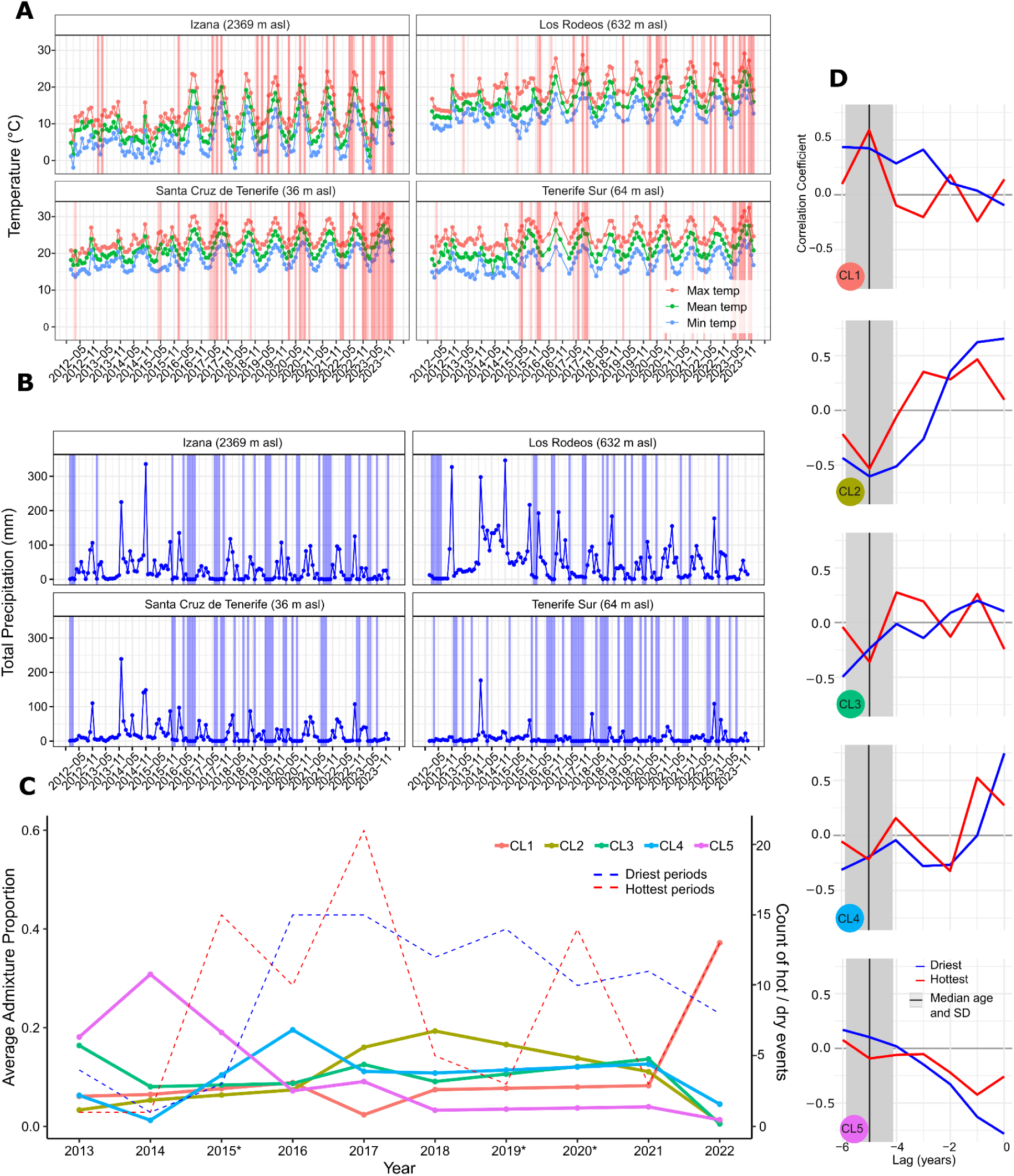
A) Maximum, mean, and minimum temperatures at monthly intervals measured at four weather stations in Tenerife between 2012 and 2023. Red bars represent mean or maximum temperatures exceeding the top 10% of temperatures in a given month. Overlapping bars from mean and maximum temperatures show as darker bars. B) Total precipitation at monthly intervals for four weather stations in Tenerife for the same timespan. Blue bars represent values below the bottom 10% of total precipitation in a given month. C) Admixture proportions of climate loci and number of extreme climate events over time. Admixture proportions interpolated in missing sampling years are indicated with a * on the x-axis. D) Cross-correlation coefficient analysis for each admixture group showing lag between climate events and changes in admixture proportions. Median estimated age and standard deviation of lizards sampled in 2022 is shown as vertical black line and grey bars (5 years, ± 0.88).

### Functional changes in promoter sequences

The kernel density plot of Tajima’s D values across promoter regions shows a high density of similar (positive) values between -700 and -1000 bp upstream of the transcription start site (TSS) and another high-density region with similar (neutral) values just before the TSS (-100 to -300 bp upstream) (Figure 7A, 7C). A small region of negative Tajima’s D values was detected between -500 and -1200 bp upstream, contrasting with the neutral loci plot, which displays only neutral Tajima’s D values around the TSS, lacking pronounced positive or negative regions (Figure 7C). The heatmap highlights that the *ETFA* promoter sequence showed the highest overall Tajima’s D score (Figure 7B), with positive values upstream of the TSS and a distinct negative region between -800 and -1000 bp. In contrast, *AQP3* displayed the lowest overall score, with negative values -400 to -1200 bp upstream. *AQP4* and *SLC25A33* also showed negative Tajima’s D values from -1400 to -1200 bp upstream. Although *HSP90B1* showed an overall low Tajima’s D score, much of its sequence approximated zero, with isolated positive and negative regions.

**Figure 7.**
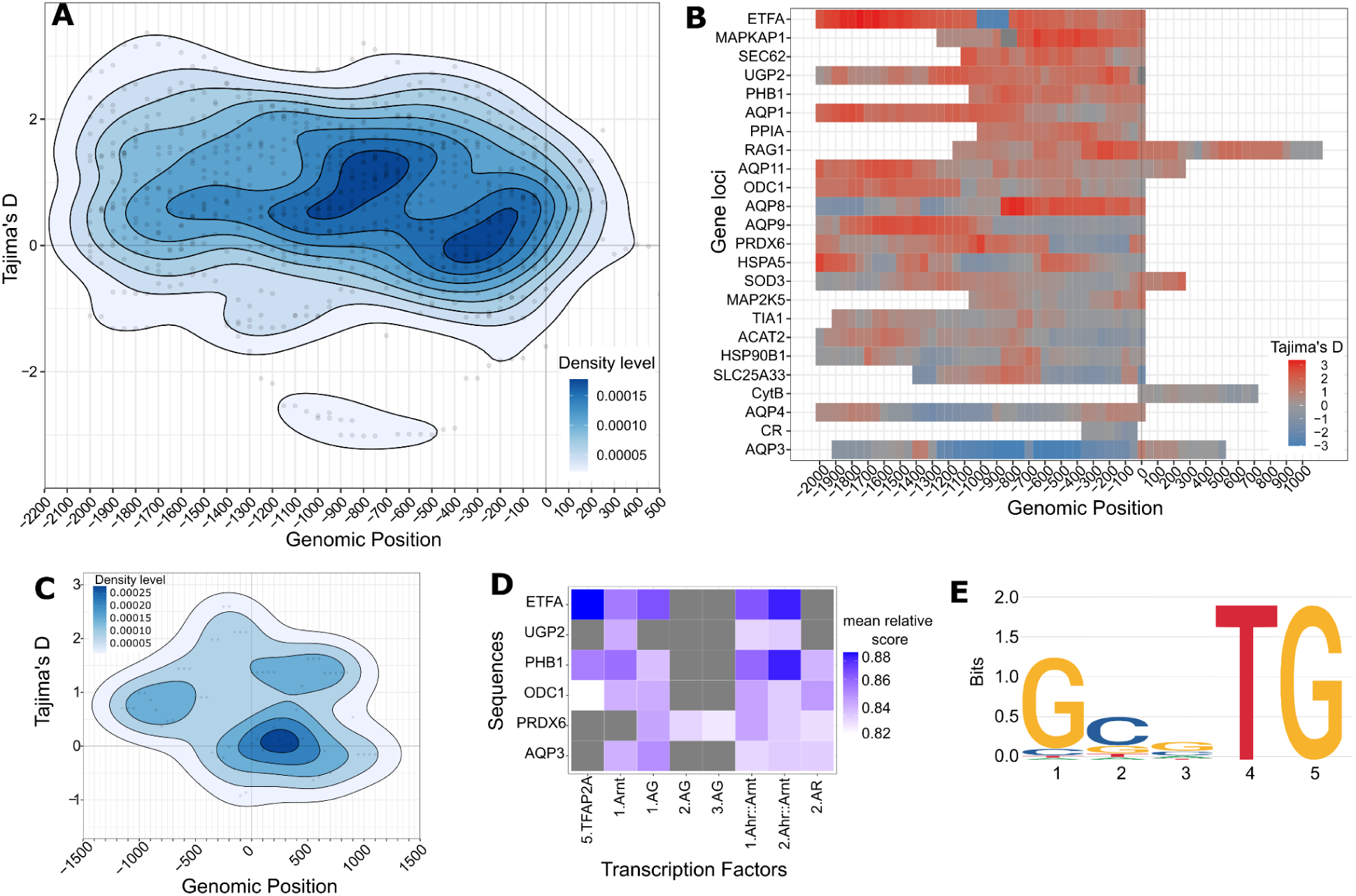
Tajima’s D across genomic positions and Transcription Factor Binding Sites. A) Kernel density plot, where darker colours represent a higher frequency of values at that position. B) shows a heatmap of Tajima’s D across genomic positions for each loci, where red colours represent positive Tajima’ D values, grey represents neutral (∼ 0), and blue represents negative. The loci are ordered from high to low in total Tajima’s D values. C) Kernel density plot just for the neutral loci (nuclear *RAG1*, mitochondrial CR and *CytB*). D) Heatmap of the mean relative scores of transcription factor binding sites for each sequence returned from JASPAR. Different motifs recognised by the same TFs are prefixed with numbers. E) Sequence motif logo demonstrating the position specific frequency of nucleotides across the sequences inputted into JASPAR for the *Ahr::Arnt* complex.

Additionally, TFBS were identified within the -700 to -1000 bp promoter region, with high (and low, for *AQP3*) Tajima’s D values. Motifs recognised by *TFAP2A* (Transcription Factor AP-2 Alpha), *Arnt* (Aryl hydrocarbon receptor nuclear translocator), *AG* (MADS box), *Ahr::Arnt* (Aryl hydrocarbon receptor::Arnt), and *AR* (Androgen Receptor) were identified with the highest relative score (Supplementary Table 8). The *ETFA* promoter had the highest relative *TFAP2A* and *Ahr::Arnt* scores, and *PHB1* had high *Ahr::Arnt* scores. The *Ahr::Arnt* motif was present in all six tested promoters. In *ETFA*, the ∼80 bp sequence surrounding the TFBS for *Ahr::Arnt* is conserved within a region of negative Tajima’s D, suggesting possible positive selection at this site (Figure 7E). This analysis does not disentangle temporal from spatial adaptation.

## Discussion

Understanding the adaptive capacity of organisms to varying environmental conditions is crucial in the context of climate change, which ultimately identifies adaptive limits and tipping points (Thurman et al. 2020; Hoffmann & Sgrò 2011). While plastic responses (e.g., behaviour or life history) offer short-term adaptation and is beneficial in effective thermoregulators (Garcia-Porta et al. 2019; Berger et al. 2021), long-term persistence requires genome-level changes including gene regulation and expression mediated by natural selection, particularly at embryonic or early developmental stages (Waldvogel, Feldmeyer, et al. 2020; Brázda et al. 2021; Young et al. 2015). These molecular adaptations underpin local adaptation and help to assess population vulnerability during the Anthropocene (Razgour et al. 2019). This study highlights the evolutionary role of promoter sequence adaptation in environmental adaptation and cellular stress tolerance at rapid temporal scales and across elevation-induced climatic gradients. This offers critical insight into how rapid genomic shifts can mediate resilience across spatially and temporally heterogeneous conditions under climate change.

### Spatial patterns of climate adaptation in promoter regions

Patterns in neutral loci align with past studies that describe the phylogeography of *G. galloti* on Tenerife, identifying two distinct clades (Thorpe, Mcgregor, and Cumming 1993; Richard and Thorpe 2001; Brown et al. 2006), and showing evidence of environment-driven divergence (Brown, Paterson, and Risse 2016). Despite the lack of samples from the southeast, the distinct patterns revealed by neutral and climate loci suggest that demographic history and environmental adaptation influence their evolutionary trajectories differently. NL revealed higher F_ST_ values from environment C corresponding to *G. g. eisentrauti* and Roque de Fuera, the offshore islet home to *G. g. insulanagae* (Martín 1985), suggesting limited gene flow. In contrast, CL displayed broader introgression across Tenerife, with five ancestry components: CL1 in the North, CL3 and CL4 at high elevations (Environment D), and CL5 in the South. Roque de Fuera de Anaga shows a distinct CL2 lineage, also shared across other regions. The alignment of these ancestry groups with distinct elevation-induced climate demonstrates the influence of environmental selection along a gradient that is decoupled from neutral genetic structuring (Hoban et al. 2016). The separation of samples into two distinct elevational bands aligns with that of the drought-induced tree line on the island (Jonsson et al. 2002), with the canopy altering the thermoregulatory landscape, and modified hydric resources between low and high elevations.

Sample-level genotype-environment associations indicate climate-related selection in distinct geographic regions. High-elevation samples from Environment D associate with annual precipitation, mean solar radiation, and inversely with mean temperature, suggesting that altitude-specific stressors, such as high radiation and low temperatures, drive selection for climate adaptation (Sun et al. 2018; Fu et al. 2022; Delgado-Suazo & Burrowes 2022; Serén et al. 2024). Conversely, genotypes from environment A in the hot, low-lying southern coast show strong associations with high temperature and low precipitation, where extreme heat may challenge activity patterns (Herrando-Pérez et al. 2019; Rutschmann et al. 2021), add to cumulative stress load (Megía-Palma et al. 2020; Megía-Palma et al. 2024), and simultaneously influence evolution through other mechanisms such as heat-driven increased nucleotide substitution rates (Garcia-Porta et al. 2019; Berger et al. 2021). While environment C did not strongly correlate with specific variables, other unmeasured factors like wind exposure and humidity likely contribute to the adaptive landscape, and local precipitation patterns may not be effectively captured in the WorldClim model such as mid-altitude North-oriented dew precipitation (Ritter et al. 2015).

The overlap of detected SNPs between the more conservative LFMM and the RDA highlights specific loci, such as “*AQP11*_1809”, as important for environmental adaptation. A notable number of SNP outliers (42%, 35/83) were identified in the *ETFA* promoter, a mitochondrial enzyme crucial for energy production and lipid metabolism (Stelzer et al. 2016; Henriques et al. 2021; Jin et al. 2021), both mechanisms which support thermal acclimation in ectotherms (Sun et al. 2022; Bickler & Buck 2007; Storey & Storey 2004). ETFs generate oxidative stress during mitochondrial energy production (Salerno et al. 2022; Henriques et al. 2021), and variants under thermal stress may have poorer performance (Henriques et al. 2011). Given the role of oxidative stress in environmental stress (Giraud-Billoud et al. 2019; Schröder & Krutmann 2005), and previous evidence that mitigation of oxidative stress helps *G. galloti* adapt across the environment (Gilbert et al. 2024), *ETFA* promoter evolution, and hence mitochondrial function, is relevant for understanding climate adaptation (Jin et al. 2021). This aligns with studies linking mitochondrial adaptations to climatic gradients in lacertids and other species under extreme cold conditions (Wollenberg Valero et al. 2022; Yudin et al. 2017).

SNPs identified in genes related to morphogenesis; regulating cell growth, organ development, and angiogenesis, may facilitate morphological and cellular adaptations to high-elevation hypoxic stress (Wollenberg Valero et al. 2022; Serén et al. 2024). At high elevations, such as on El Teide, hypoxic conditions are physiologically demanding, providing a strong selection pressure (Pamenter et al. 2020; Storz & Scott 2019). High-altitude adaptation in lizards has been explored in the Qinghai-Tibet Plateau, where hypoxia and oxidative stress adaptations support survival above 4,500 m asl (Li et al. 2023; Yang et al. 2015; Tang et al. 2013; Yang et al. 2014). In contrast, Tenerife’s smaller area (∼2000 km^2^) and 3700 m elevational range demonstrates that hypoxic adaptation can evolve within a limited geographic range and a single widespread species (Rodríguez et al. 2017), rather than through speciation across wider distributions.

Environmentally relevant SNPs in immune response genes, including those regulating cytokine activity and antiviral functions, suggest climate-induced immune modulation. Stress-related immune responses, such as through Interleukin-1 (Goshen & Yirmiya 2009), illustrate how high temperature can be immunosuppressive (Han et al. 2020) and increase parasite load (Oppliger et al. 1996; Oppliger et al. 1998; Megía-Palma et al. 2020; Megía-Palma et al. 2024), adding cumulative burdens on lizards (Tobler et al. 2015). In *G. galloti* this adds to environmental stresses from thermal constraints, water scarcity, and food availability (Rodríguez et al. 2008; Vickers et al. 2011; Dupoué et al. 2020; Megía-Palma et al. 2020).

Outlier SNPs in AQP genes involved in osmotic regulation indicate their potential role in hydric adaptation. Hydric balance in reptiles compared to temperature is equally, if not more important, in physiological adaptation across landscapes (Lillywhite 2017; Carretero et al. 2016; Rato & Carretero 2015; Carneiro, Diana, Enrique García-Muñoz, Anamarija Žagar, Panayotis Pafilis, and Miguel A. Carretero 2017; Le Galliard et al. 2021; Sannolo et al. 2020; Sannolo & Carretero 2019; Sannolo et al. 2018). AQP SNPs correlate with environmental gradients, such as low elevation, high temperature, and low precipitation, presumably supporting water stress management in arid southern locales. Studies in *G. galloti* demonstrate changes in water turnover rate during dry periods (Vernet et al. 1995) and variable instant evaporative water loss (EWL) at the landscape level (Albaladejo et al. 2022; Serén et al. 2023), indicating this is a crucial mechanism facilitating local ecophysiological adaptation, which may be explained by adaptive AQP SNPs. Future studies should further explore hydric physiological factors, such as EWL, in relation to AQP expression and variants. Heat-shock protein (HSP) SNPs, particularly in southern populations, suggest cellular defence against environmental stress via protein folding and oxidative stress mitigation (Sørensen et al. 2003). A previous study measuring GRP94 (*HSP90B1*) in *G. galloti* across the landscape linked expression to thermal, solar radiation and topological parameters, and reduced oxidative stress (Gilbert et al. 2024).

Mutations in the promoter of *HSP90B1* identified as being under balancing or purifying selection depending on the environment, add to this evidence in how HSPs may provide resilience to thermal extremes.

Outlier SNPs in *SLC25A33*, *PRDX6*, and *MAP2K5* promoters, linked to oxidative stress response, emphasise the importance of ROS regulation in broader thermal adaptability, beneficial for climate adaptation (You & Chan 2015; Sinenko et al. 2021; Ritchie & Friesen 2022). SNP associations with *ETFA* and HSP gene promoters reinforce the role of oxidative stress management in local adaptation, with plausible immune system interactions providing additional costs. Specifically, future studies should investigate oxidative damage separately for males and females, due to the large sexual dimorphism of *G. galloti* and the higher parasite load in females on Tenerife (Megía-Palma et al. 2020).

SNPs detected across environments can also be extrapolated to the landscape level, with an adaptive gradient from the high elevation interior based on SNPs in *ETFA*, *SLC25A33*, *UGP2* and *PHB1* promoters, radiating out towards coastal regions. A second adaptive gradient extends from coast to interior, involving different SNPs in AQPs, *SLC25A33*, and *ETFA* promoters, highlighting environment-dependent SNP frequency shifts within the same promoters. This genetic adaptation across elevations aligns with similar findings in other ectotherms (Bachmann & Van Buskirk 2021; Rodríguez et al. 2017; Sun et al. 2018; Yang et al. 2014). The present approach offers a landscape-scale view of adaptive genetic variation across this subtropical island, informing targeted conservation strategies for populations most vulnerable to climate change (Meek et al. 2023).

### Temporal trends in promoter climate adaptation

Temporal genetic shifts reveal climate adaptation patterns in *G. galloti* populations, with declining F_ST_ in NL over nine years suggesting reduced population differentiation likely due to increased gene flow (Reynes et al. 2024). Conversely, CL displayed increased F_ST_ in 2022, indicating rapid allele frequency shifts. This is reflected in the PCA, which shows more recent samples (2022) clustering, consistent with a reduction in allelic richness. This indicates an overall population contraction and might reflect the purging of lower-frequency alleles and loss of ancestry lineages, with one dominant set (CL1) of beneficial climate alleles increasing in relative frequency (e.g. Bourne et al. 2014).

Across all environments, allelic richness declined over time, with notable differences between interior (Environments B and D) and coastal (Environments A and C) populations, indicating temporal variation is not equivalent across habitats. Coastal populations displayed lower but more constant allelic richness, likely from more stabilised yet extreme selective pressures in the hot and dry southern environment (A) and forested northern environment (C), constraining the range of adaptive alleles (Bradshaw 1988; Cox & Cox 2015). These coastal areas also experience the greatest anthropogenic pressure, particularly from tourism, causing habitat degradation and impacting *G. galloti* populations (Otto et al. 2007; García-Romero et al. 2023). Interior populations exhibited higher allelic richness which declined over time, consistent with multivariate environmental selection pressures such as temperature extremes (Muir et al. 2014; Schweizer et al. 2021; Bodensteiner, Gangloff, et al. 2021), precipitation patterns (Llewelyn et al. 2018; Grant & Dunham 1990), oxygen pressure (Moore 2017; Jiang et al. 2021), radiative related variables (Thurman et al. 2014; Reguera et al. 2014), and other factors (Walsh & Blows 2009). Environment B, which bridges coastal and interior areas, showed the steepest allelic richness decline, and may act as a contact zone if high and low elevation populations are locally adapted (Greenbaum et al. 2014; Austin et al. 2013). A decreased trend in allelic richness could be linked to decreasing connectivity or migration (Schmidt et al. 2020; Krawiec et al. 2015;

Armas-Díaz et al. 2024), however this was not reflected in the F_ST_ values, and Tajima’s D was lower or decreased over time in environment B, indicating selective forces must play a role (Cvijović et al. 2018; Lohmueller et al. 2011). Overall, declining allelic richness over time is in this case likely to be linked to population decline, possibly due to the soft selective sweep of adaptive alleles (Messer & Petrov 2013; Hermisson & Pennings 2017).

Tracking Tajima’s D reveals different temporal selection patterns across loci: cluster 1 exhibited increasing intermediate-frequency alleles, suggesting a process of purging of rare variants in all environments except D, which lacked this pattern. In contrast, negative Tajima’s D in cluster 2 suggested purifying selection for specific alleles, likely driven by selective sweeps rather than demographic bottlenecks, as both contraction and expansion in different loci was observed (Vitti et al. 2013). This observation supports the idea that specific climate-responsive genes were targeted by selection pressures, leading to purging of maladaptive rare alleles in cluster 1 and retention of beneficial alleles in cluster 1 and 2 (Holderegger et al. 2006).

Cluster 3 and the NL displayed near-neutral Tajima’s D, suggesting stabilising selection or neutrality. With all genes contained in cluster 3 being known for their involvement in the stress response, this supports the idea of some general stress-responsive genes being evolutionarily conserved to maintain versatility against different and changing specific stressors (de Nadal et al. 2011). Constant Tajima’s D in the NL and their spatiotemporal haplogroups indicates that the selective pressures were specific to CL, and potentially, climate-induced selection events, leaving neutral regions unaffected (Holderegger et al. 2006).

As a possible explanation for the drastic shifts in ancestry group dominance observed in CL, but not NL sampled in 2022, extreme climate events, notably the hot, dry conditions of 2017 likely influenced admixture shifts over time. This aligns with the hypothesis that extreme climatic events exert selective pressures that could drive immediate adaptive responses in lizard populations through changing allele frequencies (Campbell-Staton et al. 2017; Simon et al. 2023; Wikelski & Thom 2000). The increase of CL1 in 2022 aligns with high-temperature, low-precipitation conditions from 2017, reflecting a time-lagged response where selective pressures favoured a genotype’s resilience to climate stress (Horníková et al. 2024). This time-lagged response reflects the average age of 5 years for samples collected in 2022 (estimated age of maturation 3-5 years; Serén et al. 2023), indicating that individuals are most vulnerable during early life before reproductive maturity (eggs or yearlings) (Grant et al. 2017; Vincenzi et al. 2017; Sanger et al. 2018). Adults are better able to thermoregulate and manage water loss, making hydration-sensitive thermoconforming eggs and embryos a potential bottleneck stage for climate selection. In contrast, CL2-4 showed declines, likely outcompeted by CL1 under these extreme conditions, emphasising the role of extreme weather events as a selective force in shaping population structure (Rank & Dahlhoff 2002; Jump et al. 2006).

The decreasing correlation of CL5 with lag may not be closely tied to climate extremes, conferring resilience in varying conditions, where it dominates the southern sampling sites pre-2022. The admixture patterns are indicative of selective pressures acting within a single generational event (Ehrlich et al. 2021; Ehrlich et al. 2023). This generational alignment with climatic events implies that evolution is rapidly shaping admixture proportions of selected genes in a short timescale, rather than a gradual change across multiple generations, providing a real-time glimpse into the evolutionary responses of these populations to extreme weather events.

### Functional changes in climate adaptation gene promoters

Beyond mean trends in Tajima’s D over space and time, functional motifs of adaptive significance were uncovered (Cadzow et al. 2014; Naidoo et al. 2018). Negative Tajima’s D in a small region in the *ETFA* promoter and *AQP3* points to an excess of rare alleles, indicative of recent selective sweeps. This aligns with evidence that *AQP3* evolved under positive selection in reptiles, enabling survival in drier or hotter environments (Zang et al. 2019). This means that selection favours variants that could aid *G. galloti* hatching success and drought adaptation.

Regions with negative Tajima’s D correspond with enriched transcription-factor-binding sites (TFBS), notably the “*Ahr::Arnt*” complex in the *ETFA* and *PHB1* promoters, which already demonstrate environmentally relevant phenotypic outcomes (Furue et al. 2014; Sondermann et al. 2023). *Ahr::Arnt,* consisting of the aryl hydrocarbon receptor (AHR) and its partner ARNT, regulates responses to environmental pollutants and stressors such as oxidative stress or hypoxia, and interacts with HSPs (Swanson 2002; Swedenborg & Pongratz 2010; Sondermann et al. 2023; Vorrink & Domann 2014; Cox & Miller 2004; Heid et al. 2000). This TFBS, associated with adaptive functions like immune response and melanogenesis upon UV exposure (Furue et al. 2014), likely plays a role in managing stress and facilitating adaptation in *G. galloti* (Zhang & Huang 2022). In particular, the negative Tajima’s D in *ETFA* approximately 950 bp upstream coincides with the *Ahr::Arnt* TFBS motif, suggesting selective pressure favouring variants that may alter transcriptional regulation (Kurafeiski et al. 2019). No outlier SNPs were detected in this specific region, indicating that the selection pressures on the TFBS may be more aligned to temporal changes, corresponding to the allelic richness and F_ST_ evidence for selective sweeps mentioned above, as opposed to balanced spatial heterogeneity. While experimental validation is required, the consistent appearance of the motif across multiple genes indicates selection favouring rapid regulatory adaptability, likely essential for adaptation to changing weather patterns across Tenerife.

## Conclusion

This research highlights the adaptive significance of non-coding regulatory regions in response to environmental variability in space and time, offering a molecular perspective on both long-and short-term evolutionary responses to changing climatic factors. Environmentally relevant candidate SNPs involved in oxidative stress mitigation indicate mechanistic pathways of climate adaptation, with some promoter regions exhibiting selection for high allelic diversity reflecting the environmental variability on Tenerife, and others being directionally selected by weather extremes at the temporal scale. Our findings demonstrate how regulatory sequences can facilitate rapid adaptation through a turnover in genotypes to changing conditions in the versatile lizard *G. galloti*, which could happen particularly at younger, more vulnerable life stages. This is pertinent as climate-related pressures in the Anthropocene challenge species to retain adaptive potential amidst habitat fragmentation, pollution, and extreme weather events. Our findings uncover a critical role of rapid evolution within promoter regions of relevant genes, in understanding the consequences of accelerated transformation of ecosystems.

### Funding and Acknowledgements

EG, PBA & KWV were supported by the Leeds-York-Hull Natural Environment Research Council (NERC) Doctoral Training Partnership (DTP) PANORAMA under grant NE/S007458/1. AŽ was funded by the Slovenian Research and Innovation Agency (ARIS) (P1–0255 and J1–2466). MAC acknowledges projects of MINECO (Spain) /ERDF CGL2015-67789-C2-1-P, and PGC2018-097426-B-C21, FCT (Portugal)/ ERDF 28014 02/SAICT/2017 and 2022.03361.PTDC. KWV acknowledges funding by the European Union (ERC, MolStressH2O, #101044202). Views and opinions expressed are however those of the author(s) only and do not necessarily reflect those of the European Union or the European Research Council Executive Agency. Neither the European Union nor the granting authority can be held responsible for them.

We would like to thank all members of OdysysLab (Dublin and Hull) for helpful discussions. We thank R. Donnelley and E. Chapman for assistance in the lab, and M. Wade with sequencing. We thank A. Maka for providing artwork of each subspecies, and M. Benito Smeele from ExSitu Project for photographs. We thank Acevedo-García as well as G. Albaladejo, G. Palomar, B. Fariña, J. Piquet, JL. Herrera, E. Serrano, C. Romero, X. Santos, U. Dajčman, M. Krofel, S. Novak, V. Perc and S. Reguera who assisted us in the field or shared their samples with us. We acknowledge the valuable help of IPNA-CSIC and, particularly, Manuel Nogales.

## Data availability

Complete sequence data used in the project is available on NIH NCBI, with BioProject ID: PRJNA1197140. The complete set of consensus sequences, data used, and scripts can be found in the Electronic Supplementary Materials at 10.5281/zenodo.14284296 and will be made available upon manuscript acceptance.

## Ethics and permits

Ethical approval of research for EG and MLD was obtained from the University of Hull, FEC_2022_103. The Spanish Dirección de Agricultura, Ganadería y Alimentación of Consejería de Medio Ambiente y Ordenación del Territorio of Comunidad de Madrid provided ethical approval (PROEX 128/19) after clearance of the Ethical Committee of CSIC and approval of Área de Protección Animal of Comunidad de Madrid. CAM provided a certificate for animal experimentation to RMP (0945555854754694309162) and Generalitat de Catalunya provided certificate for animal experimentation to MAC (27/07/2001). EG and MLD collection permits were obtained from Cabildo de Tenerife Sigma 1701-22 AFF 154-22. Gobierno de Canarias and Cabildo Insular de Tenerife provided specific permits AFF469/13 (2013-02234), AFF 51/16 (2016-00480) to study lizards to capture *G. galloti* across the island. This included specially protected areas such as Teide National Park and the Integral Natural Reserve of Roques de Anaga: 369/2014 (2014/2721), AFF 218/14 (2014-01172/29679), MDV/amp, AFF 110/16 2016-01215, JLRG/arm, no. 3/332, YMG/cpa, AFF 57/17 2017-00308, AFF 67/17 2017-00335, JLRG/arm (138/2). Collection permits for samples of 2021 were obtained from Cabildo de Tenerife Sigma 2021-01359 AFF 146-21 and Gobierno de Canarias Ref. Expte. 2021/26145 and for 2018 samples from Cabildo de Tenerife Sigma 2018-02258 AFF 160-18. Sampling permits AFF 57/17 2017-00308, exit record: 13413, AFF 67/17 2017-00335, exit record: 15807, AFF 160/18 2018-02258, exit record: 28466, AFF 146/21 (2021-01359), and 2021/26145; and Teide National Park sampling permit JLRG/arm, exit record 138/2 issued by Cabildo Insular de Tenerife.

## Author contribution statement

EG, KWV, and PBA contributed to the conceptualisation, investigation, data curation, formal analysis, validation, visualisation, and writing of the original draft. RMP, AMZ, MLD, NS, MAC contributed tissue samples and data. GSS assisted with lab work and Oxford Nanopore sequencing. All authors contributed to the final manuscript version.

## Conflict of Interest statement

The authors declare no conflict of interest.

## Notes

### Competing Interest Statement

The authors have declared no competing interest.

