## Supplementary Materials for "Climate drives long-term landscape and rapid short-term promoter evolution in the Western Canaries lizard, *Gallotia galloti*"

### Supplementary materials
### to the manuscript

#### “Climate drives long-term landscape and rapid short-term promoter evolution in the Western Canaries lizard, *Gallotia galloti*”

Edward Gilbert[1,2], Rodrigo Megía-Palma[3], Anamarija Žagar[4,5,6,7], Marta López-Darias[8], Miguel A. Carretero[4,5,9], Nina Serén[4 5], Graham S. Sellers[1], Pedro Beltran-Alvarez[10], Katharina C. Wollenberg Valero[11,12*]

**Contains:**
**6 Supplementary Figures**
**9 Supplementary Tables**

**Data and code availability**
Basecalled sequences are uploaded to NIH NCBI GenBank under a Sequence Read Archive (SRA) submission with BioProject ID: PRJNA1197140.
FASTA consensus sequences, the data used, and relevant code are found in the Electronic Supplementary Materials at 10.5281/zenodo.14284296, and will be made available upon manuscript acceptance.

Note: Variance partitioning of models, generation of pRDAs, and adaptive index across the landscape followed the tutorial described in (Capblancq & Forester 2021). Only minor amendments to their pipeline were made due to the context of our data, and for data visualisation.

#### Supplementary methods

##### Primer design
For each of the genomic target regions, custom primers were designed to target an upstream promoter region (from a conserved primer binding site), the 5’ Untranslated region (5’UTR), and the entire first exon (see Electronic Supplementary Materials). Genes of interest were downloaded from ENSEMBL for the lacertid lizards *Zootoca vivipara* (NCBI GenBank GCA_963506605.1), *Podarcis muralis* (NCBI GenBank GCA_004329235.1), *Lacerta agilis*

(NCBI GenBank GCA_009819535.1), including the 5'UTR and 3000 bp upstream (5' flanking sequence). These regions were mapped to the *G. galloti* genome (GCA_042478105.1) in Integrative genomics viewer (IGV; (Robinson et al. 2011)), to locate the orthologous sequence in *G. galloti*. Conserved primer binding sites in the upstream region were estimated from manually generated alignments in MEGA11 11.0.10 (Tamura et al. 2021) of the *G. galloti* genome, and the genomic sequences from other lacertids. This allowed estimates of key genomic features such the 5'UTR and the transcription start site on the *G. galloti* genome to be included in the amplicons. A respective sequence from the *G. galloti* genome was then designated as primer binding sites and primers were designed using IDT PrimerQuest, with a forward primer at the conserved upstream region, and a reverse primer at the end of the first exon (or occasionally the subsequent intron, if it was conserved). Primers were designed to maximise the size of the fragment, to include at least some protein coding information, and immediate upstream information (promoter sequences). The average target region length was 2769 bases, and ranged from 1470 bp – 6140 bp. For the longer amplicons, “internal” primers were designed to amplify the target region in two sections (Electronic Supplementary Material).

##### PCR, pooling, library preparation

Each PCR amplicon was separately amplified for each sample (236 samples * 26 amplicons = 6,136 reactions). Each reaction consisted of a total of 10 µl, described in Supplementary Table 1.

**Supplementary Table 1.** PCR reagents and volumes required for one PCR. The volumes were multiplied by the amount of products being amplified plus error, and a master mix of the Taq, DNA, and water was made.

| Ingredient | Volume x1 (µL) |
|---|---|
| LongAmp Taq 2X Master Mix | 5 |
| F+R primers (10µM) | 0.8 |
| DNA template | 0.8 |
| Nuclease free Water | 3.4 |

Thermocycling conditions were according to the Master Mix manufacturing instructions, with an annealing temperature of 55 °C, an extension time of 3:30, and final extension of 8 minutes (Supplementary Table 2).

**Supplementary Table 2.** PCR thermocycling conditions used for all amplicons, modified from manufacturers guidelines of LongAmp *Taq* 2X Master Mix (M0287, New England BioLabs).

| **Stage** | **Initial denaturation** | **Denaturation** | **Annealing** | **Extension** | **Final extension** | **Cool** |
|---|---|---|---|---|---|---|
| No. cycles | X1 | X32 | | | X1 | |
| Temp °C | 94.0 | 94.0 | 55.0 | 65.0 | 65.0 | 4.0 |
| Time (min) | 0:30 | 0:20 | 0:30 | 3:30 | 8:00 | □ |

After successful amplification, all amplicons per individual lizard were pooled in even quantities to consolidate into a single pooled sample. To ensure an even relative concentration for each amplicon in the pooled sample, the fluorescent band intensity for each amplicon as a proxy for concentration was quantified using gel electrophoresis images on ImageJ (Schneider et al. 2012), and inputted to a formula to calculate pooling quantity. The formula was calculated using a subset of samples from a prior sequencing run, and accounted for sequencing length, GC content, and band intensity. Sequencing bias was detected based on the length of the sequence and the GC content, following (Whitford et al. 2022). The formula serves to generate a value considering all these parameters so that coverage is approximate across all amplicons for one sample. The model to generate the coefficients, calculated in R, looked as follows:

*lm(depth ~ GC_percent + length + pixel_density)*

The full equation to account for the variables in the model is as follows:

$$X = -109pd - 0.05L + 23GC + 210$$

Where:

*X* = relative concentration (arbitrary units)

*pd* = pixel density (number of pixels)
*L*= sequence length (bases)
*GC* = GC content (%)

To scale the values of the equation appropriately, a min-max normalisation approach was taken. For each amplicon of each sample, the full equation was subtracted from the lowest equation value within each sample, which was divided by the maximum equation value subtracted from the minimum equation value, as follows:

$$\frac{X - min}{max - min}$$

This provided a range of 0-1, where 0 was the strongest amplicon, and 1 was the weakest amplicon. To translate into an appropriate volume to pool, these values were multiplied by 7.5 (maximum amount of PCR product available; µl). For final pipetting, values were rounded to two decimal places, and values less than 1, including zero, were rounded up to 1.

The pooled PCR products for each sample were then cleaned to remove excess PCR reagents and short oligos that may interfere with the sequencing run. This followed a solid phase reversible immobilisation (SPRI) magnetic bead capture method (adapted from (Rohland & Reich 2012)) using Sera-Mag SpeedBead Carboxylate-Modified Magnetic Particles (cytivia). The pooled amplicons were mixed with 0.6X volume of bead solution for each sample, and left to stand for 10 minutes. They were placed on a magnetic stand until the supernatant was clear, and the supernatant was discarded. The beads were incubated with a wash solution twice, allowed to dry for 5 minutes, and then the captured DNA was eluted and transferred to a fresh tube. Finally, the purified product was quantified using the Qubit HS Assay Kit.
The samples were prepared according to the Native Barcoding Kit 96 V14 (SQK-NBD114.96) protocol for amplicons provided by Oxford Nanopore Technologies. Three barcoding libraries were used and loaded on three separate MinION flow cells (FLO-MIN114). Library preparation included end-prep, native barcode ligation for each sample, adapter ligation, and clean up. Priming and loading the flow cells was done according to the manufacturer's instruction for the MinION flow cell with R10.4.1 chemistry. A total of 15 fmol of the final library was loaded for sequencing. Each sequencing run was conducted on a MinION Mk1B device through MinKNOW software (version 24.02.16) for 72 hours.

**Data processing and analysis**

WorldClim2 variables were tested for collinearity with the R package *corrplot* (Wei & Simko 2024), with variables removed when collinearity with another variable was > 0.9 (Supplementary Figure 1).

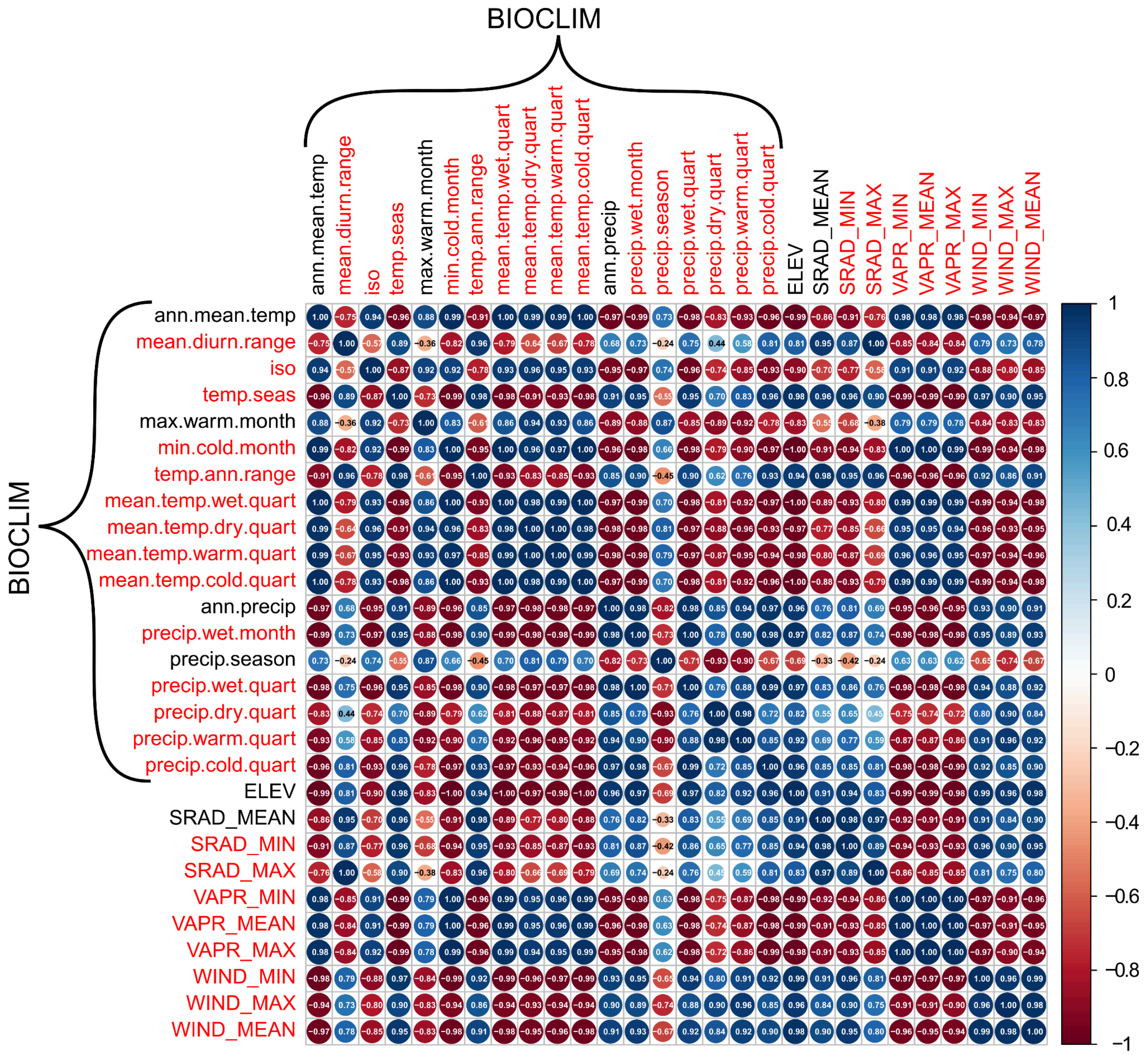


**Supplementary Figure 1.** Correlation plot of all WorldClim2 variables. Bioclimate variables are identified as “BIOCLIM” and variables retained for analyses are coloured black. Blue colours represent positive correlations and red colours represent negative correlation.

#### Spatial analysis outliers

Outliers with the furthest distance from the centroid were annotated for visualisation purposes.

An additional simple “unconstrained” model was run solely using the environmental variables (no conditioning variables accounting for population structure), as this approach can be overly conservative (Forester et al. 2018), particularly given observed higher admixture in the climate loci than the neutral loci. Given that accounting for population structuring could either hide true positives (Capblancq & Forester 2021), but not accounting for it could increase false positives (Capblancq & Forester 2021; Excoffier et al. 2009), especially given an amount of confounding variation across models (Supplementary Table 3), the degree of overlap between a simple RDA (unconstrained by population structure) and a partial RDA (constrained by population structure) was compared as in (Capblancq & Forester 2021). Outlier SNPs occurring in both the simple unconstrained RDA and the constrained pRDA were extracted, and inputted into an “adaptively enriched” RDA, following (Capblancq & Forester 2021)). This was used to create a geographical representation of adaptation across the landscape by calculating the adaptive index, using a custom function from (Capblancq & Forester 2021; Steane et al. 2014).

**Functional analysis of outliers**

To better understand the function of the genes involved, gene names from outlier SNPs shared across the GEAs were input into BioMart Ensembl online, using the Common Wall Lizard (*Podarcis muralis*) PodMur_1.0 and Green Anole (*Anolis carolinensis*) AnoCar2.0v2 datasets to extract gene ontology (GO) term attributes. GO terms were classified as either a biological process, molecular function, or cellular component, and then biological processes were further summarised by classification hierarchy. The classification of these terms are found in the Electronic Supplementary Material. The results were visualised with a chord diagram using the *circlize* package in R (Gu et al. 2014), linking each gene region where outlier loci were identified, to its general function. A weighted count table based on the number of SNPs per loci was calculated so that the chord size represents the number of SNPs.

**Temporal metrics**

Genotype information from the spatial-temporal dataset was subsetted into each major environmental category on Tenerife (see Figure 1, [A,B,C,D]), filtered for year-environment combinations with fewer than 3 samples, and the allelic richness was calculated for each of these conditions using the *allelic.richness* function in the *hierfstat* R package (Goudet & Jombart 2022). After assessing model assumptions and overdispersion, the data was fitted to a generalised linear model (GLM) with *year*env* interactions, and a gamma log link function. It was plotted as a predictive model with standard error confidence intervals. Nucleotide diversity (π) was calculated using the *nuc.div* command in the *ape* R package (Paradis & Schliep 2019) for each environment of each year. The mean nucleotide diversity across all climate loci was calculated for each environment of each year, and after checking model assumptions and overdispersion, was fitted with a GLM containing *year*env* interactions, and a gamma log link function. The model was visualised as a predictive model with standard error confidence intervals. To measure Tajima's D values at the temporal scale, *tajima.test* function from the *pegas* R package was used on the consensus fasta files subsetted yearly. Years 2013 and 2014 were removed due to small sample sizes. K-means clustering with K = 3 was used to group the genes based on temporal trends of positive, negative, and neutral, and grouped by environment type.

#### Supplementary Results

##### Sequencing

A total of 17.66 million (3.45M - 7.73M) reads were generated across all three flow cells, with 15.07 million passing quality control. After quality control, mean read count per barcode was 63,584 translating to a mean coverage of 1490 per barcode.

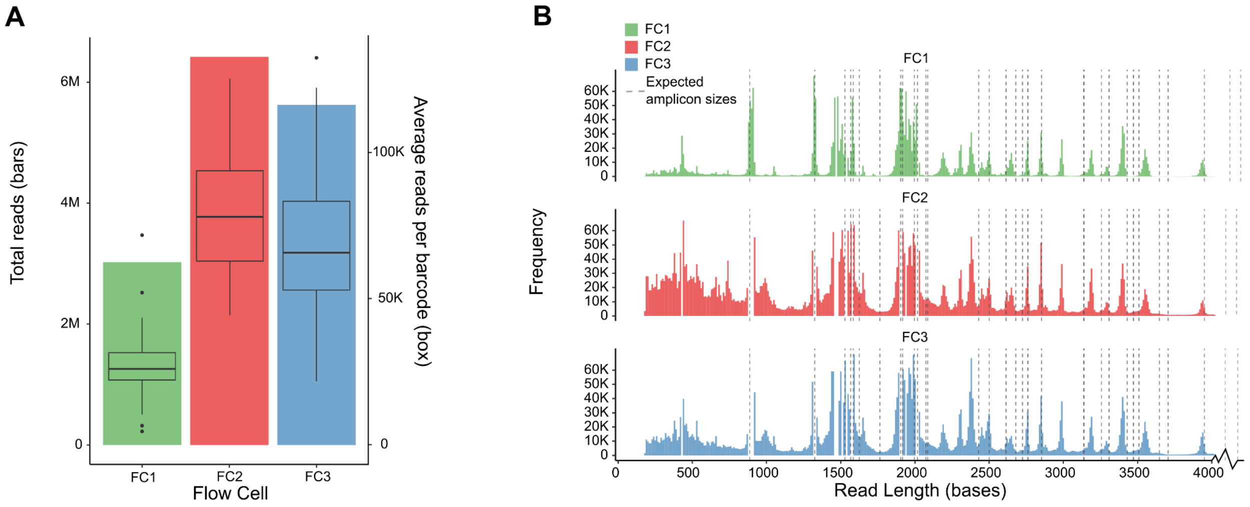


**Supplementary Figure 2.** A) Total read counts (bars) and average reads per barcode (boxplots) for each of the three flow cells. B) Read length distribution for each of the three flow cells, with dashed lines denoting the expected amplicon size. Amplicons at ~5000 bp and ~7000 bp not shown.

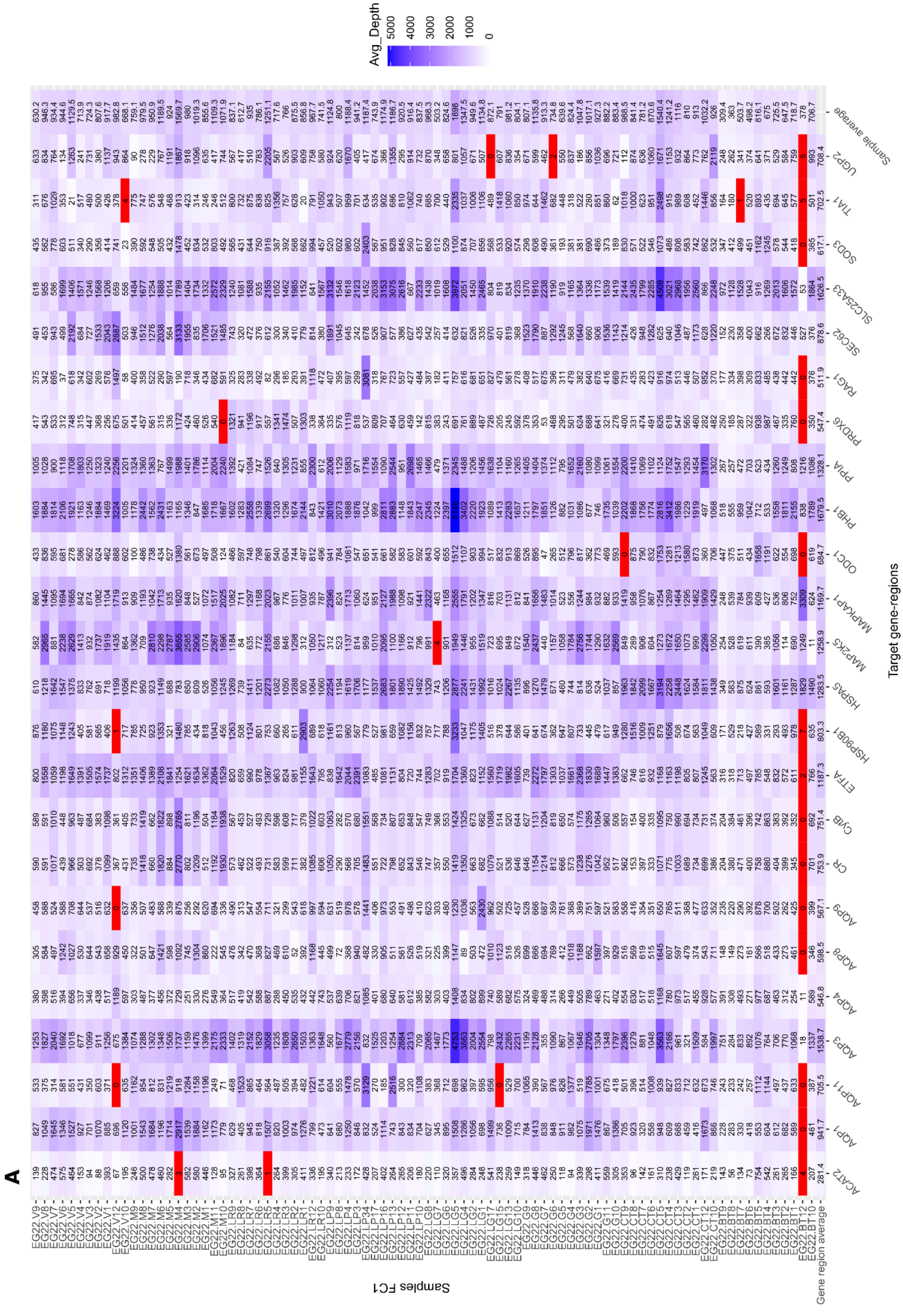
A
Avg_Depth
5000
4000
3000
2000
1000
0
Target gene-regions
Samples FC1
Gene region average
Sample average

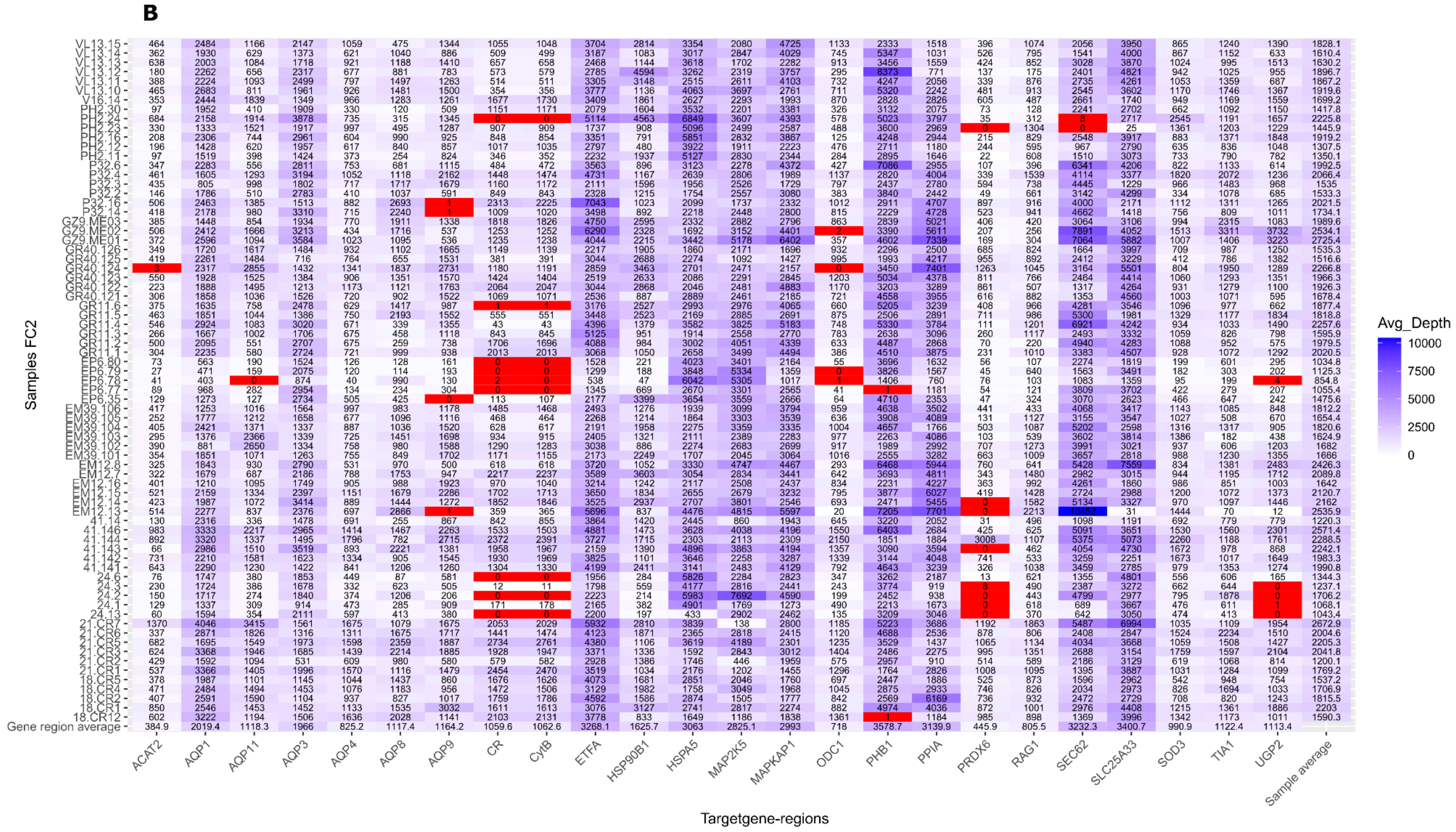
B
Samples FC2
VL13.15
VL13.14
VL13.13
VL13.12
VL13.11
VL13.10
V16.14
PH2.30
PH2.24
PH2.23
PH2.16
PH2.12
PH2.11
P32.6
P32.4
P32.3
P32.2
P32.16
P32.14
GZ9.ME03
GZ9.ME02
GZ9.ME01
GR40.126
GR40.125
GR40.124
GR40.123
GR40.122
GR40.121
GR11.6
GR11.5
GR11.4
GR11.3
GR11.2
GR11.1
EP6.80
EP6.79
EP6.78
EP6.77
EP6.35
EM39.106
EM39.105
EM39.104
EM39.103
EM39.102
EM39.101
EM12.8
EM12.7
EM12.16
EM12.15
EM12.14
EM12.13
41.14
41.146
41.144
41.143
41.142
41.141
24.6
24.3
24.2
24.1
24.13
21.CR7
21.CR6
21.CR5
21.CR3
21.CR2
21.CR1
18.CR5
18.CR4
18.CR2
18.CR1
18.CR12
Gene region average
ACAT2
AQP1
AQP11
AQP3
AQP4
AQP8
AQP9
CR
CytB
ETFA
HSP90B1
HSPA5
MAP2K5
MAPKAP1
ODC1
PHB1
PPIA
PRDX6
RAG1
SEC62
SLC25A33
SOD3
TIA1
UGP2
Sample average
Targetgene-regions
Avg_Depth
10000
7500
5000
2500
0

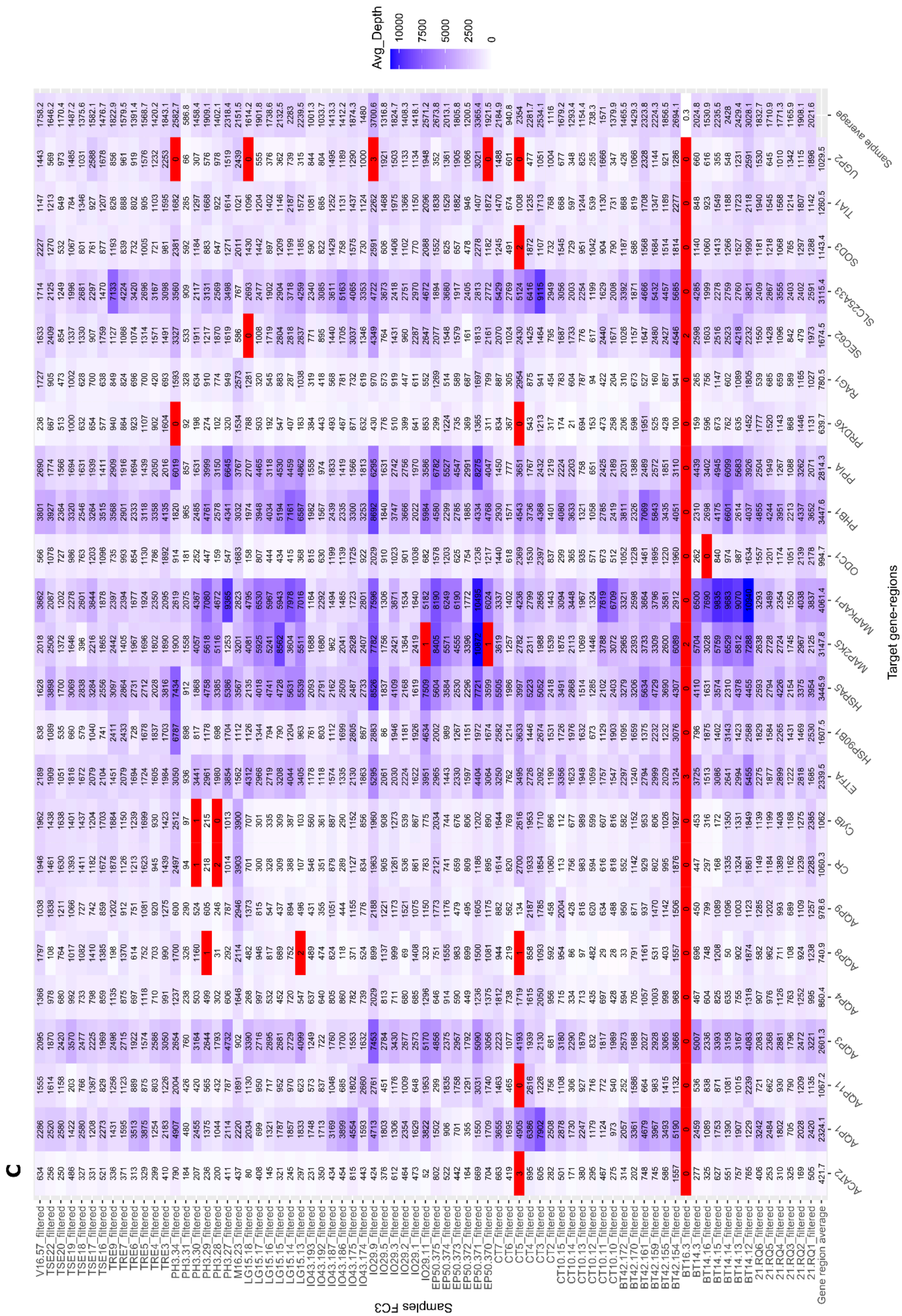
C
Avg_Depth
10000
7500
5000
2500
0
Target gene-regions
Samples FC3
ACAT2
AQP1
AQP11
AQP3
AQP4
AQP8
AQP9
CR
CYtB
ETTA
HSP90B1
HSPA5
MAP2K5
MAPKAP1
ODC1
PHB1
PPIA
PRDX6
RAG1
SEC62
SLC25A33
SOD3
TIA1
UGP2
Sample average
Gene region average

**Supplementary Figure 3.** Heatmaps demonstrating average depth for each amplicon of each sample. Each panel A, B, and C represents a separate flow cell run. Deeper colours indicate a higher sequencing depth, and red cells indicate reads < 10 for that sample sample/amplicon.

##### Variance partitioning (model comparisons)

The chosen model moving forward is the "Final" model in Supplementary Table 3, which includes the climate variables, the year and population structure principle components as conditioning variables. Supplementary Table 3 demonstrates the influence of climate, neutral genetic structure, the year, and geography on genomic variation within the dataset. In the full model, climate, population structure, geography, and year significantly explained 19% of the total genomic variance in the dataset. The effect of climate and year were highly significant when accounting for climate, population structure, and geography, explaining 3.7% and 9% of genetic variance respectively. This suggests an association between the genotype matrix (frequency of SNPs) and the environmental variables, inferring processes associated with isolation by environment (IBE). Population structuring from the putatively neutrally evolving markers accounted for just 1.8% of variance (9% of explained variation), while geographical coordinates accounted for only 1% (5% of explained variation). Although genetic structuring can be caused by both demographic processes and isolation by distance (IBD), here their contributions to our dataset was estimated independently. A large proportion of variance (28%) could not be uniquely attributed to any of the predictors included in the models, indicating potential collinearity or unaccounted for parameters. Despite this, a strong contribution to overall genetic variation of environmental parameters and temporal differences was revealed.

**Supplementary Table 3.** Comparison of partial RDA models by shifting climate, principal components, longitude and latitude, and the year, as conditioning variables. Inertia is analogous to variance, and the proportion of explainable variance demonstrates the total constrained variation explained by the "full" model.

| **Model** | **Constrained inertia** | **Model $R^2$** | **p-value** | **Proportion of explainable variance** | **Proportion variance explained** |
|---|---|---|---|---|---|
| Full | 217 | 0.09 | **0.001** | 1 | 0.19 |
| Climate | 41 | 0.01 | **0.001** | 0.18 | 0.037 |

| | | | | | |
|---|---|---|---|---|---|
| Pop. struct. | 20 | -0.0005 | 0.494 | 0.09 | 0.018 |
| Geography | 13 | -0.0007 | 0.710 | 0.05 | 0.01 |
| Year | 104 | 0.06 | **0.001** | 0.47 | 0.09 |
| Confounded climate/structure /geography/year | 40 | | | 0.28 | |
| Final | 55 | 0.02 | **0.001** | 0.25 | 0.05 |

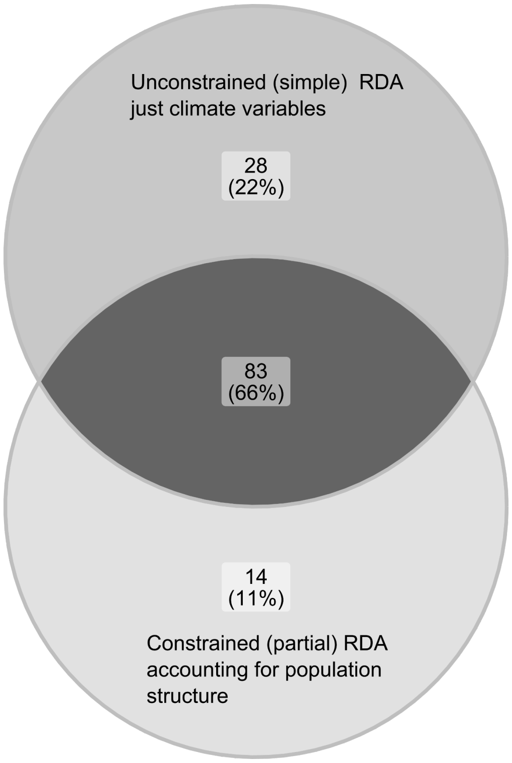


**Supplementary Figure 4.** Venn diagram showing number and proportion of SNPs detected in the unconstrained simple RDA (not accounting for population structure or year), and those in the constrained pRDA (accounting for population structure and year). 83 out of 125 (66%) of SNPs are shared between the analyses. The shared SNPs, identified as the common outliers, are used for the adaptively enriched pRDA and calculating the adaptive index across the landscape.

**Supplementary Table 4.** Generalised linear model results investigating temporal changes in allelic richness in each environment type. The model is described as *AllelicRichness ~ Year*Env, family = Gamma(link = “log”)*.

| | **Estimate** | **Std. Error** | **t-value** | **Pr(>\|t\|)** |
|---|---|---|---|---|
| Intercept (Env A) | 7.04 | 2.82 | 2.49 | 0.0128* |
| Year | -0.003 | 0.001 | -2.352 | 0.0187* |
| Env B | 10.55 | 4.54 | 2.324 | 0.0201* |
| Env C | -0.919 | 3.55 | -0.259 | 0.796 |
| Env D | 3.64 | 4.54 | 0.804 | 0.4213 |
| Year:Env B | -0.005 | 0.0022 | -2.304 | 0.0213* |
| Year:Env C | 0.0004 | 0.001 | 0.257 | 0.7973 |
| Year:Env D | -0.0017 | 0.0022 | -0.762 | 0.4459 |

**Supplementary Table 5.** Results of a Chi-squared ANOVA on the allelic richness generalised linear model.

| | **DF** | **Deviance** | **Resid. df** | **Resid. Dev.** | **Pr(>Chi)** |
|---|---|---|---|---|---|
| NULL | | | 34055 | 3607.8 | |
| Year | 1 | 1.148 | 34054 | 3606.7 | 0.0006817*** |
| Env | 3 | 209.375 | 34051 | 3397.3 | <2.2e-16*** |
| Year:Env | 3 | 0.807 | 34048 | 3396.5 | 0.04389* |

**Supplementary Table 6.** Generalised linear model results investigating temporal changes in nucleotide diversity in each environment type. The model is described as *NucleotideDiversity ~ Year*Env, family = Gamma(link = “log”).* Six values are not described due to singularities.

| | **Estimate** | **Std. Error** | **t-value** | **Pr(>\|t\|)** |
|---|---|---|---|---|
| Intercept (Env A) | -5.177 | 0.20 | -25.47 | <2e-16*** |

| | | | | |
|---|---|---|---|---|
| Year 2016 | -0.01 | 0.23 | -0.05 | 0.963 |
| Year 2017 | 0.03 | 0.23 | 0.15 | 0.883 |
| Year 2018 | 0.02 | 0.23 | 0.09 | 0.930 |
| Year 2021 | 0.06 | 0.23 | 0.28 | 0.776 |
| Year 2022 | 0.06 | 0.16 | -0.4 | 0.668 |
| Env B | -0.07 | 0.16 | -0.47 | 0.637 |
| Env C | 0.01 | 0.16 | 0.1 | 0.917 |
| Env D | -0.02 | 0.16 | 0.14 | 0.889 |
| 2016:Env B | NA | NA | NA | NA |
| 2017: Env B | 0.05 | 0.23 | 0.22 | 0.827 |
| 2018: Env B | 0.06 | 0.23 | 0.27 | 0.783 |
| 2021: Env B | 0.03 | 0.23 | 0.14 | 0.886 |
| 2022: Env B | NA | NA | NA | NA |
| 2016: Env C | 0.04 | 0.23 | 0.171 | 0.865 |
| 2017: Env C | -0.06 | 0.23 | -0.260 | 0.795 |
| 2018: Env C | NA | NA | NA | NA |
| 2021: Env C | -0.14 | 0.23 | -0.625 | 0.795 |
| 2022: Env C | NA | NA | NA | NA |
| 2016: Env D | NA | NA | NA | NA |
| 2017: Env D | -0.05 | 0.23 | -0.22 | 0.826 |
| 2018: Env D | -0.0009 | 0.23 | -0.004 | 0.997 |
| 2021: Env D | -0.06 | 0.23 | -0.278 | 0.781 |
| 2022: Env D | NA | NA | NA | NA |

**Supplementary Table 7.** Results of a Chi-squared ANOVA on the nucleotide diversity generalised linear model.

| | DF | Deviance resid. | DF resid. | Dev. | Pr(>Chi) |
|---|---|---|---|---|---|
| NULL | | | 395 | 121.29 | |
| Year | 5 | 0.599 | 390 | 120.69 | 0.8523 |
| Env | 3 | 0.092 | 387 | 120.60 | 0.9590 |
| Year:Env | 9 | 0.333 | 378 | 120.27 | 0.9992 |

**Supplementary Table 8.** Summary of output from JASPAR database highlighting higher relative scores and multiple occurrences. It summarises motifs identified from the inputted sequences.The score is calculated based on the position frequency matrix describing the binding preferences of transcription factors, comparing the likelihood of sequences occurring in the binding site vs its occurrence by chance. The relative score is a normalisation to compare across motifs of different lengths and complexities.

| Name | Score | Relative Score | Gene |
|---|---|---|---|
| Ahr::Arnt | 8.844 | 1.00 | PHB1 |
| Ahr::Arnt | 8.844 | 1.00 | ETFA |
| TFAP2A | 12.668 | 0.94 | ETFA |
| AG | 9.973 | 0.92 | ETFA |
| AR | 10.148 | 0.88 | ODC1 |
| TFAP2A | 9.445 | 0.87 | ETFA |
| Arnt | 8.609 | 0.93 | PHB1 |

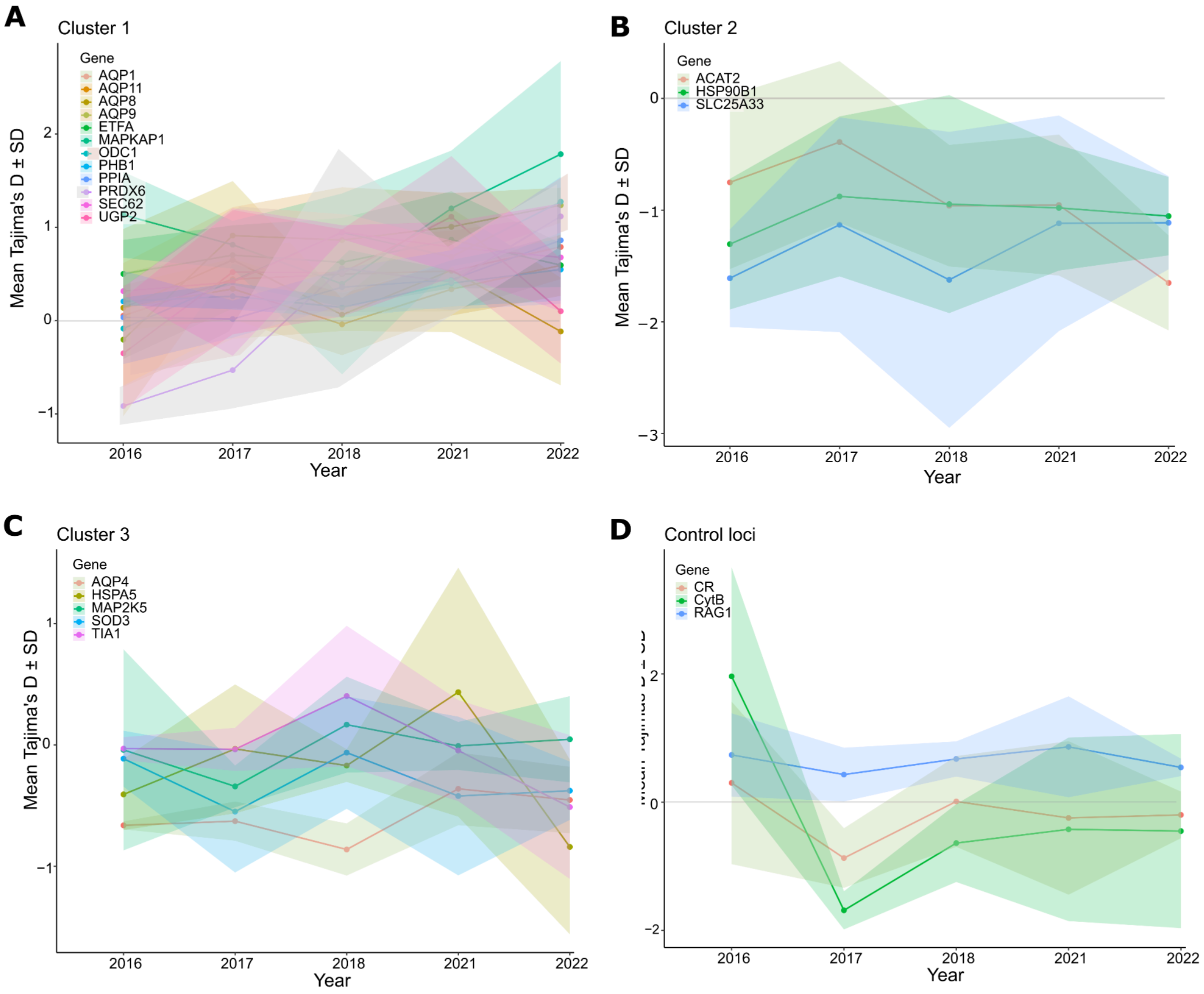


**Supplementary Figure 5.** K means clusters of mean Tajima's D values ± standard deviation (SD) from each year. Cluster 1 (A) shows genes with positive Tajima's D values (B) cluster 2 shows genes with negative Tajima's D values cluster 3 (C) shows genes with Tajima's D values approximating zero, and (D) shows the neutral loci grouped by environment type. The grey horizontal line denotes D= 0.

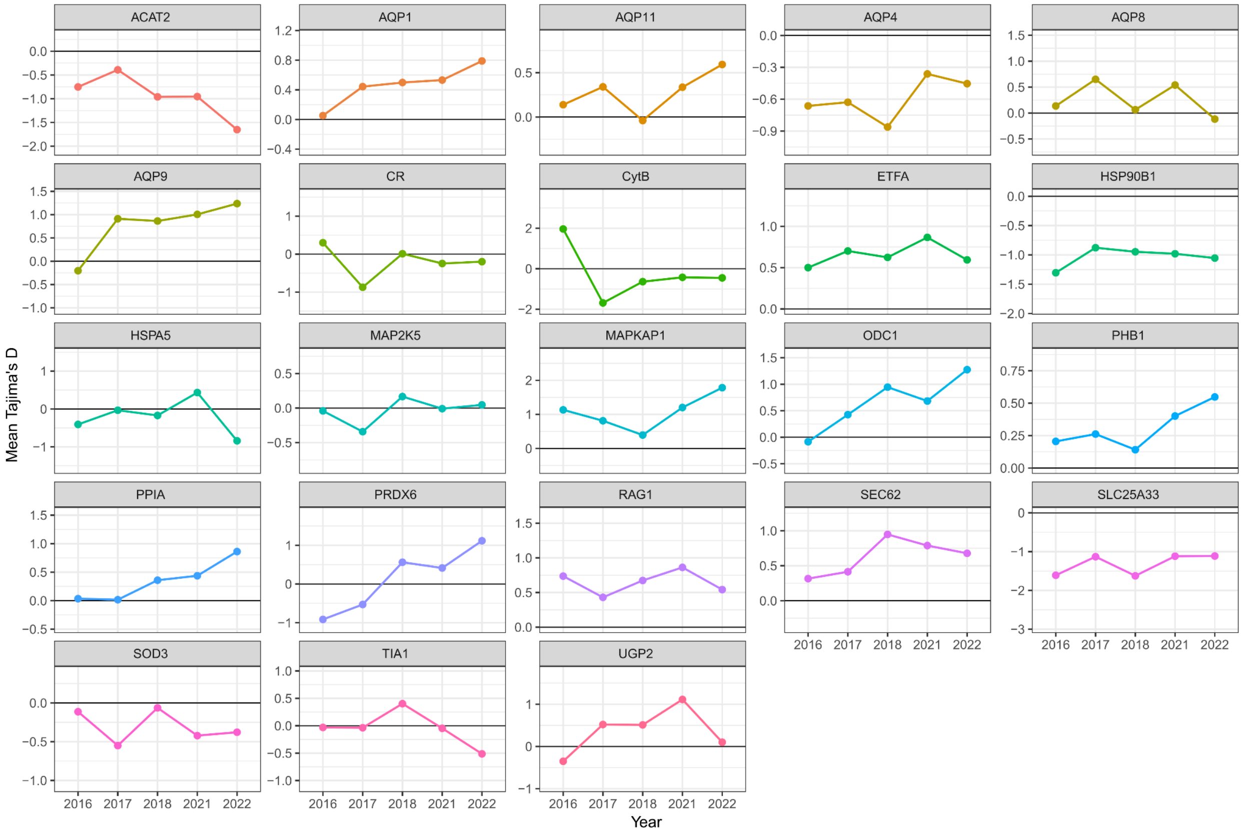


**Supplementary Figure 6.** Mean Tajima's D values over each year for each individual target gene region. Tajima's D of 0 is indicated with a black line.

**Supplementary Table 9.** Permutation testing on cross-correlation coefficients between the hottest periods, driest periods, and admixture groups over time.

| **Admixture group** | **Observed heatwave** | **Observed drought** | **Mean permutation heatwave** | **Mean permutation drought** | ***p-value* heatwave** | ***p-value* drought** |
|---|---|---|---|---|---|---|
| CL1 | 0.59 | 0.43 | 0.52 | 0.52 | 0.23 | 0.81 |
| CL2 | 0.53 | 0.65 | 0.48 | 0.49 | 0.34 | 0.12 |
| CL3 | 0.36 | 0.50 | 0.48 | 0.48 | 0.9 | 0.41 |
| CL4 | 0.52 | 0.75 | 0.53 | 0.53 | 0.5 | 0.02 |
| CL5 | 0.43 | 0.78 | 0.50 | 0.50 | 0.67 | 0.01 |
